# The central pore of HIV-1 capsomers promotes sustained stability of the viral capsid

**DOI:** 10.1101/2025.05.19.654868

**Authors:** Alex B. Kleinpeter, Donna L. Mallery, Anna Albecka, Ryan C. Burdick, Nadine Renner, J. Ole Klarhof, Boglarka Vamos, Vinay K. Pathak, Leo C. James, Eric O. Freed

## Abstract

The HIV-1 capsid, which orchestrates several key post-entry events to facilitate infection in target cells, is composed of hexamers and pentamers (capsomers) of the capsid (CA) protein arranged in a closed, conical structure known as the capsid that protects the viral RNA genome and replicative enzymes reverse transcriptase (RT) and integrase (IN). Each capsomer contains a central pore lined with rings of positively charged amino acid side chains – Arg-18 (R18) and Lys-25 (K25). The R18 and K25 rings drive capsid assembly by binding the host polyanion inositol hexakisphosphate (IP6) and are proposed to mediate the import of dNTPs into the capsid to facilitate reverse transcription. Here we demonstrate that the R18 ring can be functionally replaced by the introduction of a mutation (N21K) that establishes a new electropositive ring within the central pore. In contrast with previous studies in which R18 mutants were unable to adapt in culture, the N21K mutation facilitated the acquisition of second-site compensatory mutations that restored near-WT fitness to viral mutants lacking the R18 ring. Comparative analysis of several central pore mutants lacking the R18 ring revealed that particle infectivity was not correlated with IP6 binding or capsid assembly, but rather with capsid stability and key post-entry events including reverse transcription and nuclear entry. Our results indicate that the central pore plays critical roles in both the assembly of capsids and their sustained stability post-entry.

## Introduction

HIV-1 particle assembly occurs at the inner leaflet of the plasma membrane of infected cells. This process is driven by the Gag polyprotein precursor, which assembles with the GagPol polyprotein precursor at a Gag:GagPol ratio of ∼20:1 into a spherical lattice, called the immature Gag lattice. The oligomerization of Gag is driven by its capsid (CA) domain, and upon completion of assembly, nascent virions bud from the cell as immature virus-like particles (VLPs). Concomitant with particle release, these VLPs undergo particle maturation in which the viral protease (PR), incorporated into virions as part of GagPol, proteolytically cleaves the Gag and GagPol precursors into the mature forms of their constituent domains. The final step of productive maturation occurs when the mature CA protein assembles into a closed conical structure – called the capsid – that encloses the viral RNA genome and enzymes reverse transcriptase (RT) and integrase (IN), and serves as the protective shell of the viral core. After viral entry, the capsid remains stable and intact^1,2^, acting as a container for reverse transcription^3^, and interacting with host factors to mediate nuclear entry and viral DNA integration^4^.

The HIV-1 capsid is composed of a predominantly hexameric lattice with approximately 250 hexamers and exactly 12 pentamers of the mature CA protein. The quasi-equivalent assembly of CA into both hexamers and pentamers, collectively called capsomers, is crucial for the formation of a closed capsid during maturation. Specifically, the inclusion of pentameric “defects” at either end of the capsid lattice induces the curvature required to close the lattice, resulting in the capsid’s characteristic conical structure^5^. In the presence of high salt, recombinant CA protein assembles into helical tubes composed entirely of hexamers^6–9^. However, in the presence of the host polyanion inositol hexakisphosphate (IP6), CA assembles into capsid-like particles (CLPs) that are similar in size and shape to authentic HIV-1 capsids^10^. Indeed, IP6 is a critical co-factor for HIV-1 replication owing to its role in promoting the assembly of a stable capsid^10,11^. IP6 also promotes virus assembly by binding and stabilizing the immature Gag lattice in a process that enriches virions with levels of IP6 necessary to build the capsid during maturation^10,12,13^.

IP6 promotes the assembly of the HIV-1 capsid by binding to two amino acid side chains in the CA N-terminal domain – Arg-18 (R18) and Lys-25 (K25) – that form positively charged rings within a “central pore” in capsomers. Both the R18 ring and K25 ring can bind an individual IP6 molecule, and by neutralizing these positive charges lining the central pore, IP6 stabilizes capsomers. Mutations of either of these IP6-binding residues prevents proper capsid assembly and severely reduces particle infectivity^14–20^. Consistent with the ability of IP6 to drive the in vitro assembly of CLPs, IP6 has been proposed to play a particularly important role in the formation of pentamers^14–16,21,22^. Mutations at K25 prevent IP6-mediated CLP assembly *in vitro*, resulting instead in the assembly of helical CA tubes exclusively^14–16^. Furthermore, our previous work demonstrated that K25A CLP assembly *in vitro* and closed capsid assembly in virions is restored by second-site compensatory mutations that likely restore pentamer formation^14^. In addition to binding IP6, the central pore also binds dNTPs in an interaction that is driven primarily by R18, suggesting that dNTPs could be transported through the central pore to facilitate reverse transcription inside closed capsids^17^.

Optimal stability of the HIV-1 capsid after viral entry is critical for several post-entry functions required for productive infection. Efficient reverse transcription of the viral genome requires a closed, stable capsid^23^ to protect the viral nucleic acid from detection by host innate immune sensors^24–27^ and to keep the RT enzyme associated with its RNA substrate. Additionally, stable capsids are required for the core to efficiently dock at the nuclear envelope and pass through the nuclear pore complex (NPC)^28,29^. After nuclear entry and the completion of reverse transcription, the core undergoes uncoating, a process that is characterized by the disassembly of the capsid leading to the release of the viral DNA for integration into host chromatin^1,2,30,31^. The timing of uncoating is crucial for productive infection, and premature uncoating due to mutations that destabilize the capsid prevents efficient reverse transcription, nuclear entry, and integration^32,33^. IP6 stabilizes pre-formed capsids *in vitro*^23^, however its role in stabilizing capsids in cells is less certain. Several studies have shown that depletion of IP6 in target cells does not disrupt HIV-1 infection^13,34,35^. In contrast, the infectivity of select CA mutants with hypostable capsids is acutely reduced by IP6 depletion, suggesting that IP6 stabilizes capsids post-entry but that this stabilizing effect is not required for WT HIV-1 to complete a productive infection. IP6 has also been proposed to play a role in dNTP import through the central pore of capsomers^16^. Ultimately, while the importance of the central pore and its interaction with IP6 is clear regarding capsid assembly, the significance of this interaction in infected cells post-entry and its implications for capsid stability, dNTP import, and reverse transcription are not well understood.

Here we examine the role the central pore of HIV-1 capsomers by determining the consequences of replacing the R18 ring with a new positively charged ring in the central pore. We report that replacing the R18 ring facilitates the acquisition of second-site compensatory mutations that potently restore viral replication. Further manipulation of the central pore and comparative analysis of several mutants lacking the R18 ring reveal that the ability to bind IP6 and assemble capsids in vitro and in virions is not correlated with infectivity. However, we find that non-infectious central pore mutants lacking the R18 ring are defective for reverse transcription and nuclear entry, and that the mechanism underlying these defects is a lack of sustained capsid stability. Our findings indicate that, in addition to its well-established role in capsid assembly, the central pore plays an important part in maintaining the stability of capsids post-entry.

## Results

### The R18 ring in the central pore of capsid hexamers can be replaced by a new positively charged ring at position 21

We recently reported that propagation of an HIV-1 mutant lacking the K25 ring (K25A) resulted in the rapid selection of second-site compensatory mutations that restore the ability of HIV-1 to assemble closed capsids^14^. However, our attempts to propagate several R18 mutants did not result in the acquisition of compensatory mutations^14^. Importantly, in vitro assembly assays indicated that R18 and K25 play separate roles in capsid assembly with R18 driving CA hexamer formation and K25 driving CA pentamer formation. Therefore, we investigated how HIV-1 could adapt to a loss of the R18 ring. In an additional in-vitro selection K25A acquired the N21K substitution (Figure S1). Interestingly, N21 is located in helix 1 of CA ∼1 turn of the helix below R18 and above K25, and its side chain, like those of R18 and K25, is oriented toward the center of the central pore in the context of assembled capsomers (Fig 1A). We reasoned that the introduction of a lysine at position 21 could establish an additional positively charged ring capable of interacting with IP6, and may facilitate enhanced adaptability in HIV-1 mutants lacking an electropositive ring at amino acid 18 in CA.

**Figure 1.**
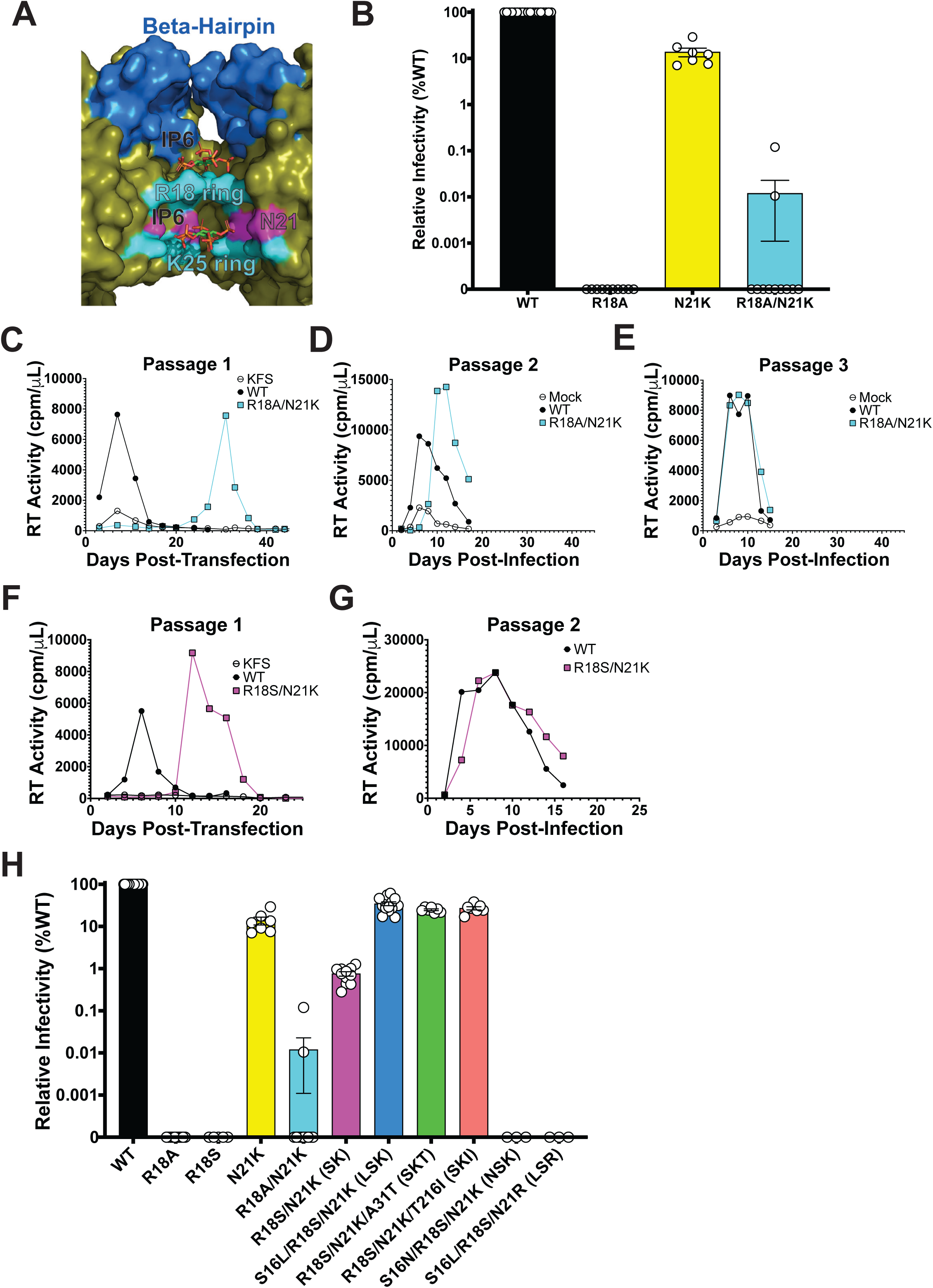
The N21K substitution facilitates the acquisition of compensatory mutations by R18A. (A) Cross section of the central pore of the CA hexamer (based on PDB 6R6Q) with IP6 molecules bound to the R18 and K25 rings. The location of N21 on the inner surface of the central pore is also shown. (B) Single-cycle infectivity of WT HIV-1 and the indicated mutants quantified by luminescence in infected TZM-bl cells. Virus stocks were normalized by RT assay prior to infection of TZM-bl cells. Error bars depict the mean ± s.e.m. from at least 7 independent experiments. (C-E) Representative forced evolution experiment depicting spreading infection of WT and R18A/N21K after initial transfection of MT4 cells (C) and a second (D) and third (E) passage of virus collected from the preceding passage. (F-G) Representative forced evolution experiment depicting spreading infection of WT and R18S/N21K after initial transfection of MT4 cells (F) and a subsequent passage (G) of virus collected at the peak of replication. The Env(-) clone pNL4-3/KFS is included as a negative control. (H) Single-cycle infectivity of WT HIV-1 and the indicated mutants quantified by luminescence in infected TZM-bl cells. Virus stocks were normalized by RT assay prior to infection of TZM-bl cells. The N21K and R18A/N21K data presented for comparison in (H) is the same that is presented in (B). Error bars depict the mean ± s.e.m. from at least 3 independent experiments. Abbreviations for the mutants used throughout the study are indicated in parentheses in (H) and listed here: R18S/N21K (SK); S16L/R18S/N21K (LSK); R18S/N21K/A31T (SKT); R18S/N21K/T216I (SKI); S16N/R18S/N21K (NSK); S16L/R18S/N21R (LSR).

To test the hypothesis that the R18 ring can be functionally replaced, we transfected 293T cells with infectious molecular clones harboring R18A, N21K, or R18A/N21K mutations to produce virions and measured their specific infectivity in TZM-bl cells. R18A displayed no measurable levels of infection in our assay, whereas N21K demonstrated a ∼10-fold decrease in infectivity relative to wild-type (WT). Like R18A, R18A/N21K infectivity was often not measurable, however we were able to detect low levels of infection in some experimental replicates, suggesting that the infectivity of the R18A/N21K mutant is near the detection limit of our assay. Ultimately, R18A/N21K demonstrated a ∼4-log decrease in infectivity relative to WT HIV-1 (Figure 1B). While these results indicate that the R18A/N21K mutant is severely defective, we postulated that this low-level infectivity may facilitate the acquisition of compensatory mutations upon propagation in T-cell lines. To test this, we transfected MT4 cells with pNL4-3-R18A/N21K and monitored replication kinetics by quantifying the RT activity in the supernatants. We observed a substantial (24-42 day) delay in replication of R18A/N21K relative to WT suggestive of the acquisition of compensatory mutations. We subsequently re-passaged virus collected from the peak of replication until replication kinetics resembled those of WT virus (Figure 1 C-E). We then extracted genomic DNA from infected cells collected at the peak of replication, PCR-amplified the Gag-coding region of the proviral DNA, and subjected these amplicons to sequencing analysis to identify compensatory mutations. We performed these selection experiments several times with multiple replicates and observed the acquisition of several potential compensatory mutations by R18A/N21K (Table S1). Two mutations were located in amino acid residues within or adjacent to the central pore, including an alanine-to-serine substitution at position 18 (A18S), S16L, and H12Y (Figure S2A). We also observed the A31T substitution which is present on the luminal surface of the CA hexamers and pentamers that comprise the capsid (Figure S2B). In subsequent experiments, we propagated R18S/N21K in MT4 cells (Figure 1 F, G) and observed a rapid acquisition of A31T or T216I, which we previously reported as a compensatory mutation that restores CA pentamer formation and closed capsid assembly to K25A^14^. These findings indicate that HIV-1 can acquire mutations both proximal and distal to the central pore capable of restoring replication to a mutant lacking the R18 ring.

To test the ability of the selected mutations to compensate for the loss of the R18 ring in the presence of N21K, we introduced each mutation into infectious molecular clones harboring R18A/N21K or R18S/N21K substitutions and compared their specific infectivity. We found that the addition of S16L, T216I, or the combination of H12Y and A31T (which were acquired together in an individual experiment, Table S1) resulted in a 65-300-fold increase in R18A/N21K particle infectivity, though these mutants remained substantially (25-100-fold) less infectious than WT NL4-3 (Figure S3A). Remarkably, the change from an alanine (R18A/N21K) to a serine (R18S/N21K) resulted in a similar increase in particle infectivity (Figure 1H). Furthermore, S16L (S16L/R18S/N21K), A31T (R18S/N21K/A31T), or T216I (R18S/N21K/T216I) restored R18S/N21K infectivity ∼30-fold to ∼30% that of WT (Figure 1H). While these mutations do not confer a complete rescue, they confer a ∼3000-fold increase in infectivity over the R18A/N21K mutant that was used to initiate the selection experiments. H12Y did not increase R18S/N21K infectivity and did not enhance the ability of S16L or A31T to restore infectivity to R18S/N21K. Furthermore, the simultaneous addition of S16L and A31T to R18S/N21K antagonized the restorative effect conferred by each mutation individually (Figure S3B). Finally, we engineered two additional mutants for comparison with S16**L**/R18**S**/N21**K** (hereafter called “LSK”). S16**N**/R18**S**/N21**K** (hereafter called “NSK”) was engineered because S16N has been previously reported to restore replication and infectivity to the replication-defective central pore mutant I15V^36^. S16**L**/R18**S**/N21**R** (LSR) was engineered to test the effect of a different positively charged residue at position 21 in the absence of the R18 ring. Interestingly, despite only differing from the rescued LSK mutant by one amino acid, each of these mutants is completely non-infectious. Finally, we tested the replicative capacity of each of these mutant viruses in MT4 cells. The results were consistent with our infectivity measurements. Briefly, over 25 days in culture, WT, N21K, LSK, R18S/N21K/A31T (SKT), and R18S/N21K/T216I (SKI) each replicated with WT or near-WT kinetics, peaking at Day 4-8 post-transfection. Consistent with selection experiments, R18S/N21K (SK) replicated with a ∼10 day delay relative to WT, whereas R18A, R18S, R18A/N21K, NSK, and LSR showed no signs of replication (Figure S4). We also measured viral replication in peripheral blood mononuclear cells (PBMCs) isolated from four individual donors (Fig S5). Whereas WT displayed rapid replication, peaking at day 4 post infection, LSK, SKT, and SKI showed delayed replication kinetics and virus production, peaking between day 8 and 10. We did not observe any signs of SK replication in PBMCs. Overall, these infectivity and replication data demonstrate while HIV-1 can adapt to the loss of the R18 ring, it remains extremely sensitive to changes in the central pore of capsomers.

### Inositol phosphates bind and stabilize capsid hexamers lacking the R18 ring

Upon confirming that the N21K substitution restores infectivity to HIV-1 mutants lacking a positively charged ring at position 18 in CA, we sought to understand how these changes to the central pore affect the ability of ligands IP6 and dNTPs to bind and stabilize CA hexamers. To address this question, we expressed and purified CA hexamers for three variants demonstrating vast differences in particle infectivity – WT (100% infectivity), LSK (∼30% infectivity), and NSK (non-infectious) – and subjected these hexamers to several biochemical and structural analyses. Our finding that N21K restores infectivity and replication to HIV-1 mutants lacking a positively charged amino acid at position 18 in CA suggests that this change functionally replaces the R18 ring with a new positively charged ring of amino acids at position 21 (K21 ring). To confirm the presence of a new electropositive ring at K21, we solved crystal structures of LSK and NSK crosslinked CA hexamers crystallized in the presence of IP6 and compared their central pores to that of our previously solved WT structure (PDB 6R6Q) in which two IP5 molecules were resolved within the central pore – one bound above the R18 ring and another coordinated by the surrounding K25 rings. While the structures of the central pores of WT, LSK, and NSK CA hexamers were largely similar (Figure 2A), the data confirmed that the N21K mutation introduces a new ring of positively charged residues just above the K25 ring. In the LSK and NSK structures, this K21 ring participates in electrostatic interactions with a molecule of IP6 bound at the center of the K25 ring. Most notably, LSK and NSK CA hexamers lack an IP6 molecule bound at the top of the pore where IP6 has been shown to interact with the R18 ring in WT CA hexamers and pentamers. Presumably, this is because the R18S mutation removes the positively charged side chains required to coordinate an IP6 molecule. Interestingly, the sharing of a single molecule of IP6 by the K21 and K25 rings in the LSK and NSK structures is reminiscent of how IP6 binds to the immature Gag lattice, in which one IP6 molecule is coordinated by the CA-K158 and CA-K227 rings at the center of Gag hexamers^10^. The similarity observed in the structure of the central pore of LSK and NSK CA hexamers and their similar interactions with IP6 suggests that the ability to bind IP6 is not the mechanism underlying the observed differences between the infectivity of LSK and NSK.

**Figure 2.**
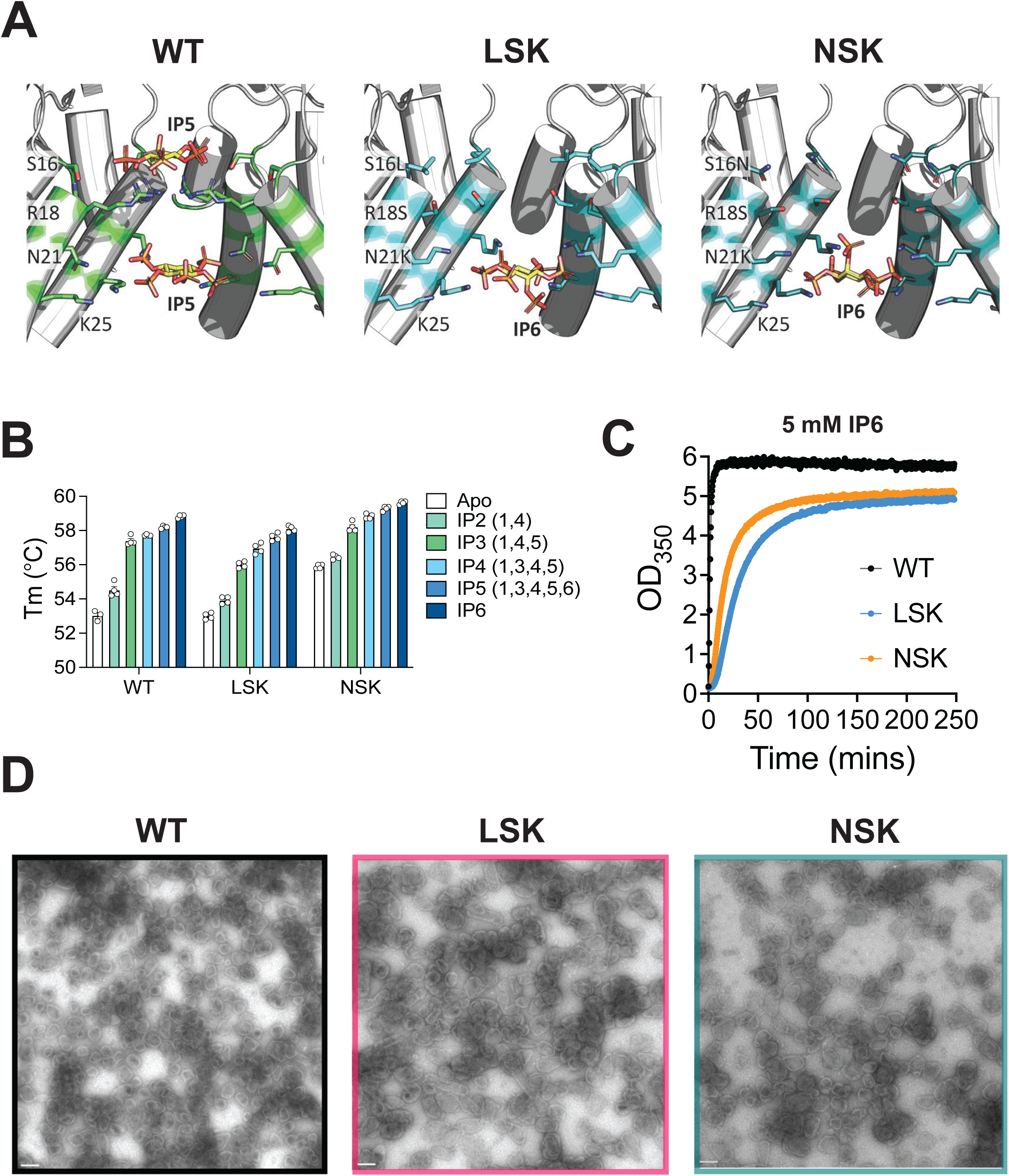
IP6 binds, stabilizes and promotes the assembly of LSK and NSK CA. (A) Cross sections of the central pore of crystal structures of LSK, and NSK crosslinked CA hexamers bound to IP6 solved in this study compared to that of the previously solved crystal structure of the WT crosslinked CA hexamer bound to IP5 (PDB 6R6Q). (B) Thermal stability measurements for the indicated crosslinked CA hexamers alone or in the presence of the indicated inositol phosphates, as measured by differential scanning fluorimetry. Error bars depict the mean (s.e.m. from at least 3 independent experiments. (C) In vitro assembly kinetics of 100 (M of the indicated recombinant mature CA protein in the presence of 5 mM IP6, as determined by measuring absorbance at 350 nm over time. (D) Negative stain EM images of the assembly reactions performed in (C). Scale bars 100 nm. Abbreviations: S16L/R18S/N21K (LSK); S16N/R18S/N21K (NSK).

To more directly compare the ability of LSK and NSK to bind IP6, we used nano differential scanning fluorimetry to quantify the extent to which IP6 and other inositol phosphates enhance the thermal stability of CA hexamers. In the absence of inositol phosphates, we observed no difference in the stability of WT and LSK CA hexamers, whereas NSK CA hexamers were significantly more stable (Figure 2B, compare white bars). Thermal stability of CA hexamers was increased by the addition of the inositol phosphates IP2 (1,4), IP3 (1,4,5), IP4 (1,3,4,5), IP5 (1,3,4,5,6), and IP6 for each variant. As expected, CA hexamers were most potently stabilized by IP6. Peak thermal stability in the presence of IP6 was highest for NSK CA hexamers, though the shift in thermal stability induced by IP6 was greater for WT and LSK CA hexamers (Figure 2B, compare white and dark blue bars). This observation is likely due to the increased inherent stability of NSK CA hexamers, resulting in a lower thermal stability shift in the presence of IP6. These data are consistent with our structural observations, which did not reveal substantial differences in the ability of LSK and NSK to interact with IP6 in the context of CA hexamers.

### NSK assembles capsids in vitro and in virions

While the use of crosslinked hexamers is useful for the study of capsid-host interactions in vitro, we sought to investigate the effect of LSK and NSK on IP6 binding in a more biologically relevant in vitro system. We therefore used an in vitro turbidity assay in combination with negative-stain electron microscopy (EM) to assess the ability of IP6 to drive the assembly of LSK and NSK CA protein into CLPs that broadly recapitulate the size and shape of authentic HIV-1 capsids. We mixed mature WT, LSK, or NSK CA protein with 5 mM IP6 and measured the kinetics of assembly by quantifying the increase in optical density at 350 nm (OD_350_) (Figure 2C). While LSK and NSK CA protein assembled with slower kinetics than WT (with NSK assembling slightly faster than LSK), negative-stain EM revealed an abundance of CLP structures present across all samples (Figure 2D). Similar results were obtained when assembly reactions were performed in the presence of 1.5 and 3 mM IP6 (Figure S6). This finding suggests that although LSK and NSK CA assembles less efficiently than WT in the presence of IP6, each can assemble *in vitro* into the closed conical capsid structure that is required for infection.

Next, we sought to determine the effect of the LSK and NSK mutations on the ability of HIV-1 to properly assemble capsids in virions. We produced non-infectious WT, LSK, and NSK virions in 293T cells and assessed capsid morphology using cryo-electron tomography (cryo-ET). For each virus, we observed a range of capsid morphologies and classified each imaged virion into one of four morphological categories: immature, mature (conical), mature (tubular), and mature (irregular) (Figure 3 A and B). Roughly 75% of WT virions displayed the conical or tubular capsid morphologies that are associated with productive infection, whereas only ∼50% of LSK and NSK virions contained conical or tubular capsids. Notably, the number of LSK and NSK virions containing irregular capsid morphologies was doubled that of WT virions. These findings are consistent with our *in vitro* assembly results showing that LSK and NSK CA can assemble into CLPs, but do so less efficiently than WT. Finally, the similarity in the distribution of capsid morphologies found in NSK and LSK virions suggests that, despite the complete lack of particle infectivity displayed by NSK, it is not defective for capsid assembly.

**Figure 3.**
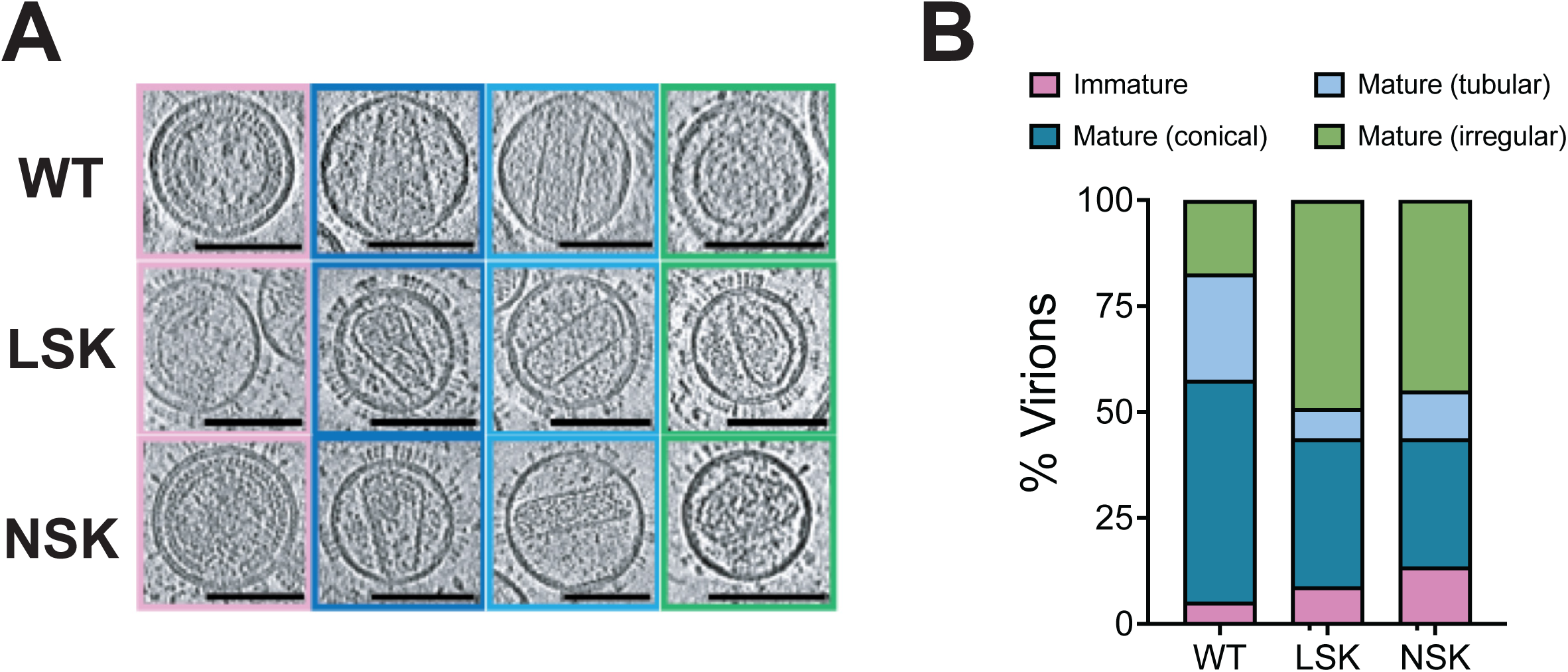
LSK and NSK assemble capsids morphologically similar to those of WT in virions. VSV-G pseudotyped WT, LSK, and NSK virions were purified and subjected to cryo-ET analysis. Tilt series were collected and reconstructions performed to assess capsid morphology. At least 100 virions for each were analyzed and classified as displaying immature morphology or conical, tubular, or irregular mature capsid morphology. Examples of sliced tomograms for each category are shown for each virus population analyzed (A) and the overall distribution for each virus population is shown in (B). Scale bars, 100 nm. Abbreviations: S16L/R18S/N21K (LSK); S16N/R18S/N21K (NSK).

### NSK is defective for reverse transcription in cells and in vitro

The central pore of capsomers and its interaction with IP6 is crucial for the assembly of the HIV-1 capsid. However, our comparison of the LSK and NSK central pore mutants, in which we deliberately manipulated the IP6 binding site, reveals that the drastic differences in the particle infectivity of these mutants cannot be explained by differences in the ability of IP6 to bind and stabilize CA hexamers or facilitate capsid assembly. To understand the mechanism underlying the differences in particle infectivity between LSK and NSK, we quantified the accumulation of RT products in SupT1 cells infected with replication-incompetent WT, LSK, and NSK virions pseudotyped with the vesicular stomatitis virus G (VSV-G) protein. We also included the R18S/N21K (SK) mutant and the rescued R18S/N21K/A31T (SKT) and R18S/N21K/T216I (SKI) mutants in this analysis for comparison. WT early (minus-strand strong stop) and late (second-strand transfer) RT products accumulated rapidly during the first 6 hours post-infection (hpi) and continued to rise slowly between 6 and 12 hpi (Figure 4 A and B). The rescued mutants LSK, SKT, and SKI showed similar reverse transcription kinetics, though the total copy numbers were ∼1.5-3-fold lower than WT at each time point. In contrast, SK and NSK displayed only minor accumulation of early and late RT products over the course of infection. Thus, the levels of SK and NSK RT products decreased progressively relative to WT at each time point collected. At 12 hpi, SK early and late RT products were reduced 20-and 50-fold, respectively, relative to WT, and NSK early and late RT products were reduced 50- and 200-fold. We also quantified the accumulation of 2-LTR circles, which form only in the nucleus and thus provide a measure of nuclear import of viral DNA. Similar to the results observed for RT products, LSK, SKT, and SKI displayed only modestly reduced levels of 2-LTR circles, whereas SK and NSK displayed drastic reductions (50-300-fold) (Figure 4C). Importantly, the reductions in RT products and 2-LTR circles for each of these mutants relative to WT correlate well with the infectivity defects displayed by each of these mutants (Figure 1H): WT > LSK, SKT, & SKI > SK > NSK. This suggests that the primary biological mechanism underlying the infectivity defects induced by replacing the R18 ring with the K21 ring is decreased efficiency of reverse transcription and nuclear entry.

**Figure 4.**
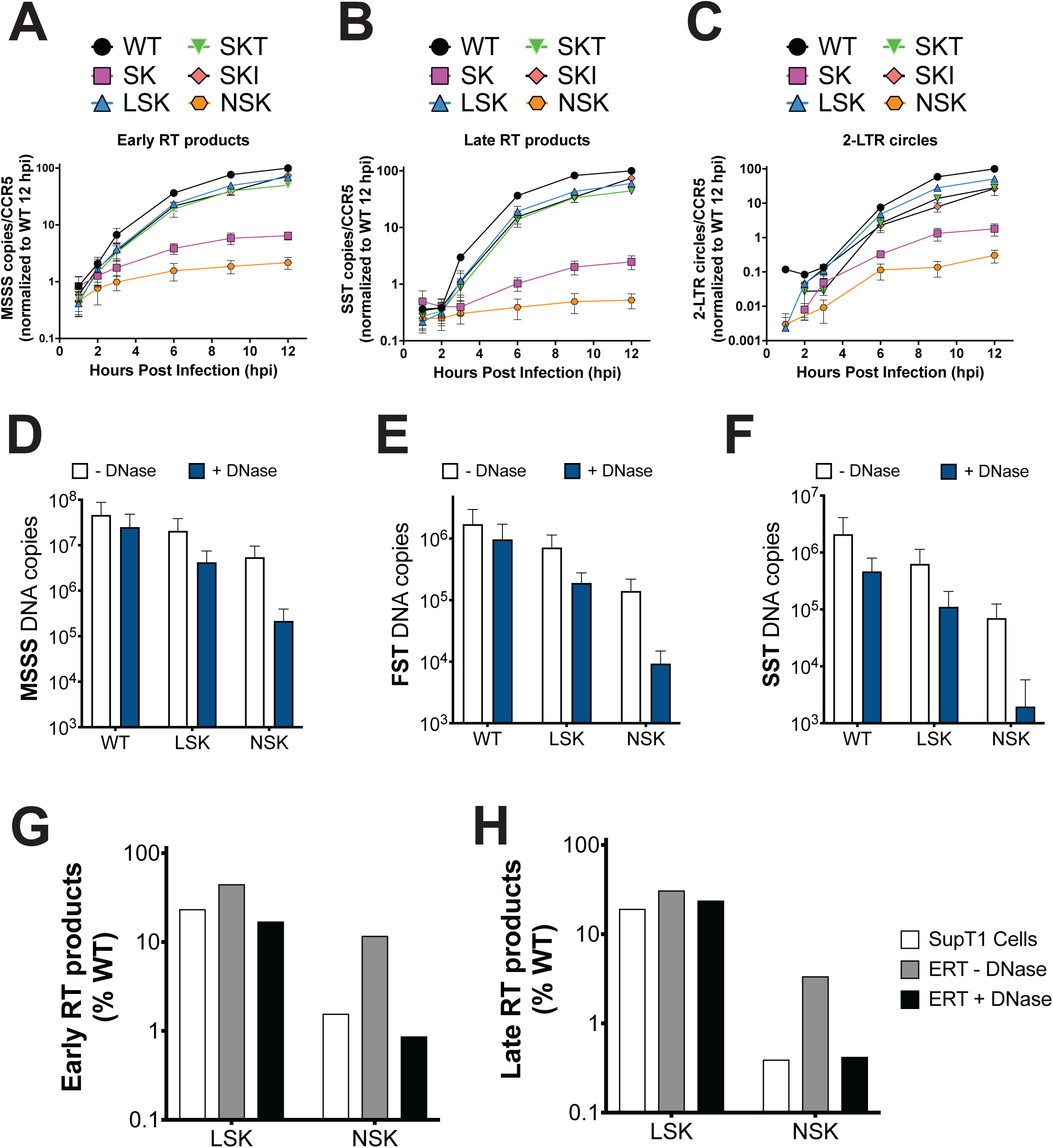
NSK virions are defective for reverse transcription in cells and in vitro. (A-C) Early RT products (A), late RT products (B), and 2-LTR circles (C) were quantified for WT, SK, LSK, SKT, SKI, and NSK at 1, 2, 3, 6, 9, and 12 hours post-infection (hpi) of SupT1 cells by qPCR. Each individual sample was quantified relative to CCR5 copy number to control for the cell number, and all data for each viral DNA species were normalized to copy number/CCR5 for WT at 12 hpi. Error bars depict the mean ± s.e.m. from 3 independent experiments. (D-F) minus-strand strong stop (D, MSSS), first-strand transfer (E, FST), and second-strand transfer (F, SST) DNA copies were quantified by qPCR in endogenous reverse transcription (ERT) reactions performed for 6 hours at 37°C with Streptolysin O (SLO) and in the presence or absence of DNase. Error bars depict the mean ± s.e.m. from at least 2 independent experiments. (G-H) Comparison of mean LSK and NSK early (G) and late (H) RT products at 6 hours post infection in SupT1 cells relative to WT to MSSS (G) and SST (H) RT products in ERT reactions performed in the presence and absence of DNase. Abbreviations: R18S/N21K (SK); S16L/R18S/N21K (LSK); R18S/N21K/A31T (SKT); R18S/N21K/T216I (SKI); S16N/R18S/N21K (NSK).

There are two possible mechanisms that may explain how the changes that we made in the central pore could affect the efficiency of reverse transcription. First, we have previously shown that R18 plays a key role in the binding to dNTPs, which have been proposed to translocate through the central pore to facilitate reverse transcription inside closed capsids^14,17^. Therefore, the replacement of the R18 ring with a K21 ring may prevent efficient dNTP binding and import into the lumen of the viral capsid, depriving RT of the substrates that it requires for viral DNA synthesis. We therefore measured the ability of ATP and dATP to stabilize WT, LSK, and NSK crosslinked CA hexamers. Whereas WT CA hexamers display increased thermal stability upon the addition of ATP and dATP, LSK and NSK CA hexamers were only marginally stabilized (Figure S7A). Consistent with this finding, fluorescence polarization (FP) assays revealed that the affinity of ATP for LSK (K_d_= 161.5 μM) and NSK (K_d_= 112.9 μM) CA hexamers is ∼10-fold lower than its affinity for WT (K_d_= 12.31 μM) CA hexamers (Figure S7B). We previously reported that the R18G mutation results in a ∼1000-fold decrease in the affinity of ATP for CA hexamers, suggesting that the introduction of the N21K mutation significantly restores nucleotide binding in the absence of the R18 ring, though not to fully WT levels. These data suggest that less efficient binding to dNTPs may contribute to the reverse transcription defects that we observed. However, the finding that the reduction in ATP binding is similar for LSK and NSK indicates that nucleotide binding to the central pore does not fully explain the vast differences in infectivity and reverse transcription exhibited by LSK and NSK.

The second mechanism that may explain how the changes to the central pore may alter the efficiency of reverse transcription is perturbed capsid stability post-entry. While the initial assembly of the capsid in virions is required for efficient reverse transcription and infection, capsids also must remain closed and stable after viral fusion, acting as a container for reverse transcription and a barrier to sensing by the innate immune system. We therefore tested the ability of WT, LSK, and NSK virions to reverse transcribe their genomes in an endogenous reverse transcription (ERT) assay. Briefly, virions were incubated with dNTPs, IP6, and streptolysin O (SLO), a pore-forming toxin that disrupts the integrity of the viral membrane, for 6 hours at 37°C. The pore-forming activity of SLO allows IP6 and dNTPs access to the interior of the virion to facilitate reverse transcription inside closed IP6-stabilized capsids. Consistent with our quantification of RT products in infected cells, the production of early (MSSS), intermediate (first-strand transfer, FST), and late RT products was slightly reduced for LSK relative to WT. However, the reduction in RT products exhibited by NSK relative to WT and LSK was considerably less severe than in cells (Figure 4 D-F, white bars; G-H, compare white and grey bars). Identical ERT reactions performed in the presence of DNase reduced the quantity of all RT products for each virus. Because DNase is too large to enter a closed capsid, we interpreted this reduction in RT products as a consequence of DNase-mediated digestion of reverse transcripts that were not protected by a closed capsid. WT RT products were least susceptible to DNase treatment, whereas LSK RT products displayed modest susceptibility to DNase. NSK RT products, particularly late RT products were highly susceptible to DNase digestion, suggesting that a large percentage of NSK RT products were not protected within closed capsids (Figure 4 D-F, blue bars). Importantly, the relative production of WT, LSK, and NSK RT products observed in the presence of DNase closely recapitulates the results we obtained in cells at 6 hpi (Figure 4 G & H, compare white and black bars). Together, our reverse transcription measurements in cells and in vitro suggest that NSK capsids are inherently unstable, leading to a loss of integrity upon disruption of the viral membrane. In cells, we posit that this loss of NSK capsid integrity occurs shortly after viral fusion and leads to the rapid disassembly of the capsid and severely reduced reverse transcription due to the diffusion of the RT enzyme from its RNA substrate. In our ERT assay, we suspect that in the absence of DNase, the loss of NSK capsid integrity occurs after permeabilization of the viral membrane by SLO. However, because the overall architecture of the viral membrane remains intact, large-scale capsid disassembly and the loss of the RT enzyme is slowed, allowing reverse transcription to continue even after loss of capsid integrity. Finally, the milder sensitivity to DNase displayed by LSK suggests that LSK capsids are considerably more stable than NSK capsids, but less stable than WT capsids.

### Mutants lacking the R18 ring demonstrate variable capsid stability, nuclear envelope docking, and nuclear entry

To test the inherent stability of the central pore mutant capsids after viral membrane permeabilization, we utilized a previously described^14^ TIRF microscopy assay (Figure 5A and B). Briefly, WT and several central pore mutants (R18S, N21K, SK, LSK, and NSK) virions were produced using a GagPol construct with EGFP inserted between its MA and CA domains and flanked by viral protease cleavage sites. During viral maturation, EGFP is passively incorporated into capsids and serves as a fluid-phase marker that is released upon capsid rupture. EGFP-labeled virions were immobilized, permeabilized with SLO in the presence or absence of IP6 for 30 minutes, and imaged to quantify the number of virions containing intact capsids. Consistent with previous results^14^, in the absence of IP6, we observed very low EGFP+ particle counts for WT and all central pore mutants relative to virus-only controls, suggesting that in the absence of added IP6 capsids rapidly rupture after membrane permeabilization (Figure 5C). In the presence of IP6, we observed markedly increased EGFP+ particle counts for WT, N21K, and LSK virions, suggesting that these capsids remained intact in the presence of IP6. In contrast, we observed substantially smaller increases in intact SK and NSK capsids in the presence of IP6, while R18S virions displayed no increase in intact capsids in this condition. These results are broadly consistent with the differences in infectivity and reverse transcription measurements for these mutants and suggest that the inherent stability of capsids is the primary mechanism underlying these observed differences.

**Figure 5.**
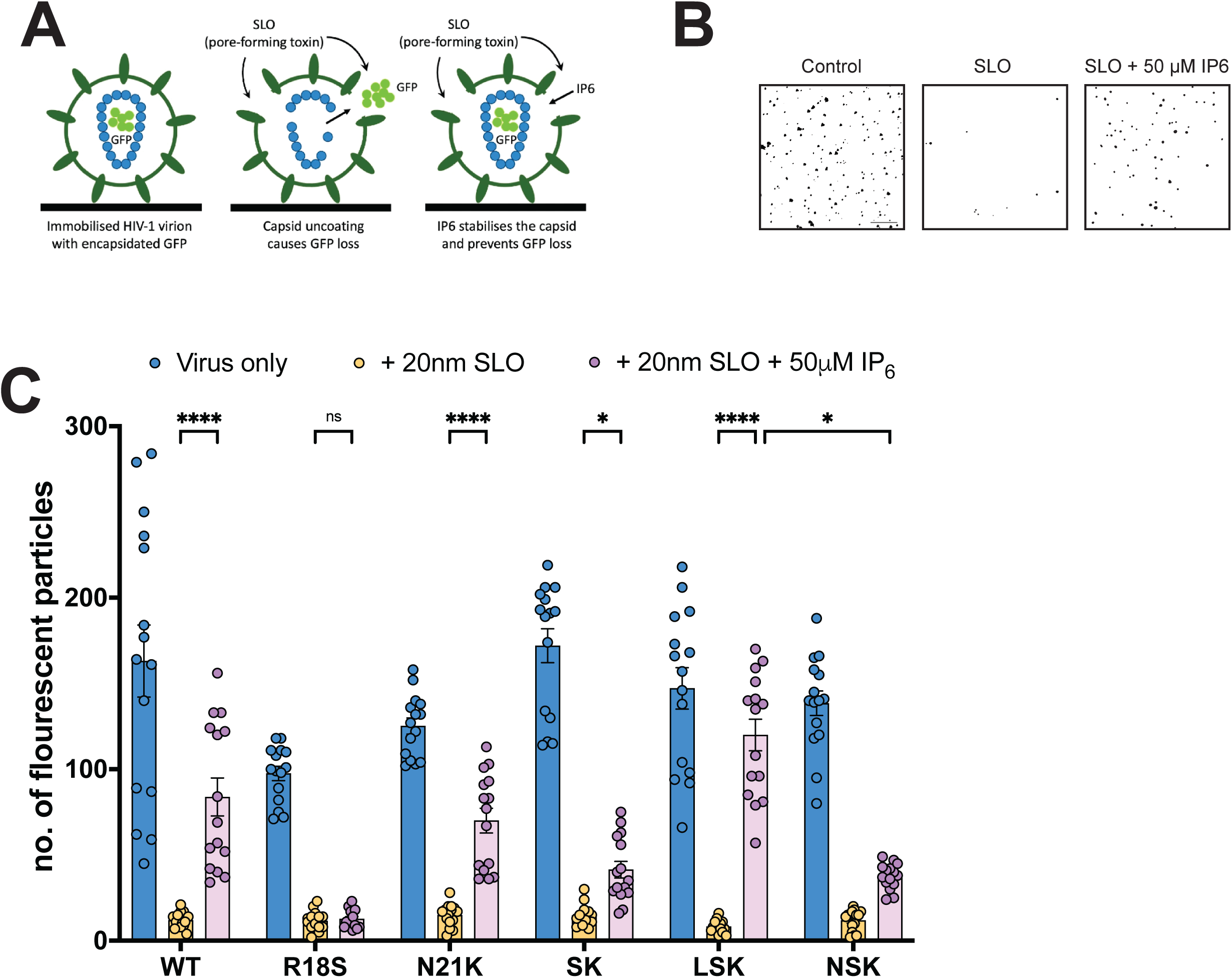
HIV-1 central pore mutants display variable capsid stability. (A) Indicated viruses containing EGFP as a capsid content marker were bound to glass dishes and permeabilized with SLO in the presence or absence of 50 μM IP6. (B) Images were acquired with a TIRF microscope and particles counted using Fiji. Representative masks generated during analysis of immobilized WT virus 30 min post-SLO or control treatment ± 50 μM IP6 are shown. Scale bar = 10 µm (C) Intact capsids were counted for the indicated mutants in untreated, and SLO-treated ± 50 μM IP6 conditions. Error bars depict the mean ± s.e.m. from counts acquired from 15 total images spanning 3 independent experiments. Statistical significance was determined by Two-way ANOVA with Dunnet’s multiple comparisons test (p-value summary: >0.05 = not significant; <0.05 = *; <0.01 = **; <0.001 = ***; <0.0001 = ****). Abbreviations: R18S/N21K (SK); S16L/R18S/N21K (LSK); S16N/R18S/N21K (NSK).

To further investigate the effect of IP6 on the capsid stability of CA mutants lacking the R18 ring, we measured the effect of IP6 depletion in target cells on particle infectivity. Several studies have reported that while WT HIV-1 infectivity is unaffected by the depletion of IP6 in target cells^13,34,35^, the infectivity of certain CA mutants that assemble inherently unstable capsids is severely reduced in such cells^35^. We therefore compared the infectivity of WT and several central pore mutants (N21K, SK, LSK, SKT, SKI) in wild-type 293T cells (parental) and cells in which IPPK, the host enzyme that phosphorylates IP5 to synthesize IP6, was knocked out (IPPK KO). As a positive control we also measured the effect of IP6 depletion on the hypostable CA mutant virus P38A. Consistent with previous reports^13,34,35,37^, WT HIV-1 infectivity was similar in parental and IPPK KO cells, whereas P38A infectivity was reduced ∼30-fold in IPPK KO cells relative to parental cells (Figure S8). Our central pore mutants showed much milder sensitivity to target cell IP6 depletion. SK and LSK infectivity was reduced ∼3-fold, and N21K and SKT displayed a 2-fold reduction in infectivity in IPPK KO cells. We also observed a statistically significant, but minor, increase in SKI infectivity in IPPK KO cells. We were unable to test NSK in these experiments as its infectivity was below the detection limit of our assay in both cell lines. The mild sensitivity to IP6 depletion displayed by LSK may be indicative of a slight reduction in capsid stability and is consistent with the mild DNase sensitivity observed in our ERT experiments. However, the SK mutant, which demonstrates significantly reduced capsid stability in our TIRF assay, displays the same mild sensitivity to target cell IP6 depletion. This suggests that the effects of target cell IP6 depletion on the infectivity of CA mutants are not necessarily correlated with capsid stability and should be interpreted with caution.

We next sought to determine the ability of our central pore mutant capsids to dock at the nuclear pore and enter the nucleus. We produced WT, LSK, and NSK virions from 293T cells expressing YFP-labeled APOBEC3F that is resistant to Vif-mediated degradation (A3F-E289K-YFP) and infected HeLa cells with these virions. A3F-YFP is a host restriction factor that is packaged into virions and is incorporated into HIV-1 capsids via its interaction with the viral ribonucleoprotein, providing a marker for the subcellular localization of capsids in infected cells^29^. Infected cells were fixed 6 hours post-infection and discrete YFP-positive particles were quantified in the nucleus and at the nuclear envelope (NE) using confocal microscopy. The percentage of NSK particles at the NE and in the nucleus was severely reduced relative to WT and comparable to the level of WT particles in the presence of the capsid inhibitor PF74, which destabilizes the capsid and prevents NE docking and nuclear entry^1^ (Figure 6). LSK capsids docked at the NE with WT-efficiency, but displayed a modest 2-fold decrease in nuclear entry relative to WT. These data are consistent with the reduction in 2-LTR circles that we observed for NSK and LSK. Based on the preceding results, and because optimal capsid stability is required for NE docking and nuclear entry^28,29^, we propose that NSK capsids are inherently unstable, resulting in essentially no NE docking, nuclear entry, or infection. In contrast, LSK capsids are stable enough to dock at the nuclear pore complex (NPC), but do not pass through the NPC as efficiently as WT capsids, leading to a modest decrease in infectivity.

**Figure 6.**
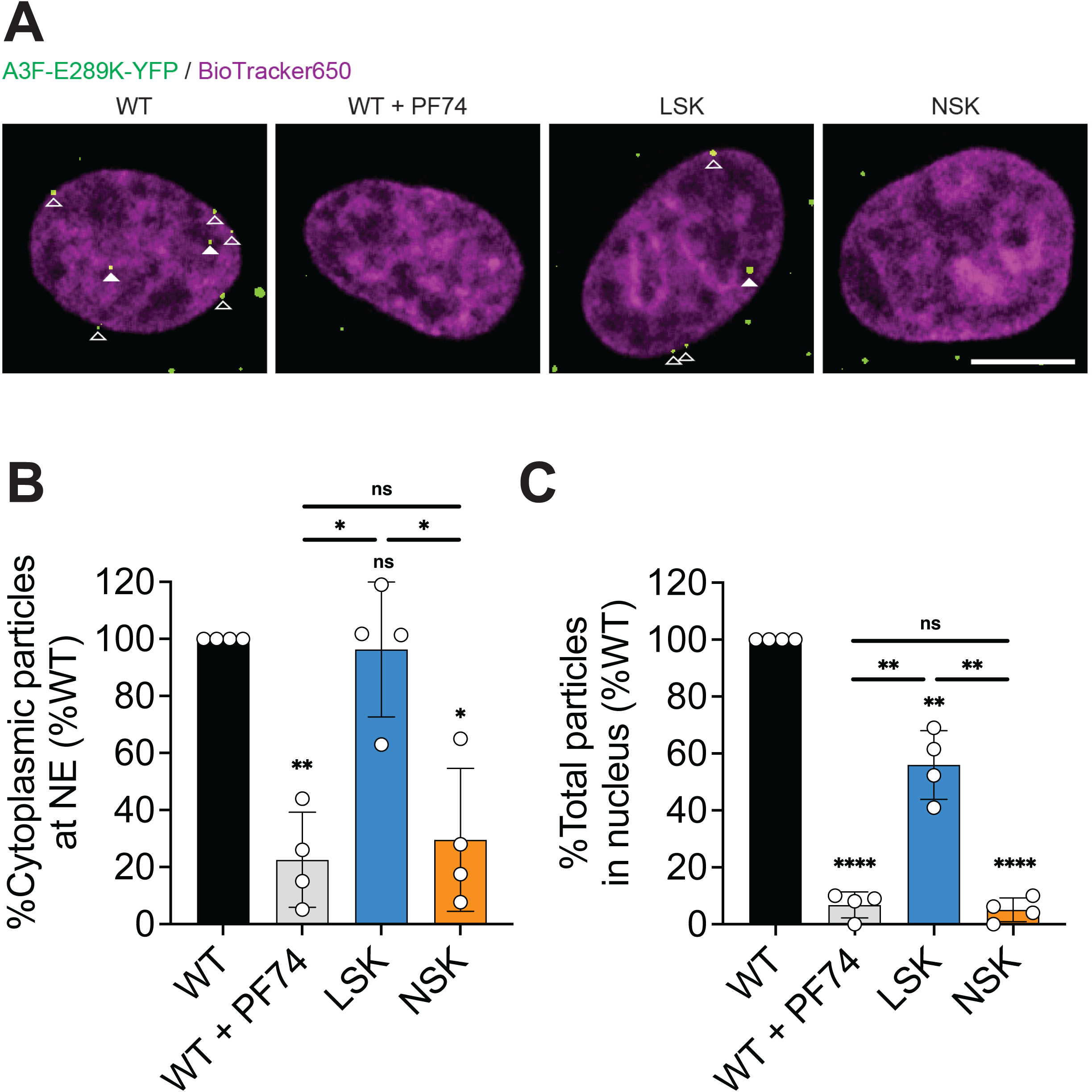
NSK capsids are defective for nuclear envelope docking and nuclear entry. VSV-pseudotyped ι1env A3F-E289K-YFP-labeled WT, LSK, and NSK virions were used to infect HeLa cells for 6 hours. Infected cells were then fixed, stained with BioTracker 650 Red Nuclear Dye, and imaged by confocal microscopy to determine the subcellular localization of HIV-1 capsids. An additional condition in which WT-infected cells were treated with 10 μM PF74 was used as a negative control. (A) Representative confocal images showing HIV-1 capsids labeled with A3F-E289K-YFP at the nuclear envelope (open triangle) and inside the nucleus (filled triangle). Scale bar, 10 μm. (B) YFP-positive particles at the nuclear envelope were quantified as a percentage of the total particles in the cytoplasm and expressed as a percentage of WT in untreated cells. (B) YFP-positive particles in the nucleus were quantified as a percentage of the total particles in the cell and expressed as a percentage of WT in untreated cells. Statistical significance was determined by paired t-test (p-value summary: >0.05 = not significant; <0.05 = *; <0.01 = **; <0.001 = ***; <0.0001 = ****). Abbreviations: S16L/R18S/N21K (LSK); S16N/R18S/N21K (NSK).

## Discussion

Each capsomer in the HIV-1 capsid lattice contains a central pore that binds IP6 to facilitate capsid assembly^10^. The central pore also binds dNTPs, which are proposed to translocate through the pore to facilitate DNA synthesis inside the capsid^16,17^. These binding events are coordinated by two positively charged rings (R18 and K25) that line the inner rim of the central pore. Our previous work produced two findings that are relevant to this study. First, we found that HIV-1 is unable to adapt to the loss of its R18 ring, whereas mutation of the K25 ring can be readily compensated for by mutations in CA that do not re-establish a positively charged ring in the central pore. Second, we found that the K25 ring is particularly important for the assembly of closed capsids during maturation. This finding is consistent with several other studies that have shown that mutations within the central pore prevent proper capsid assembly^18,32,38–40^.

In this study we engineered a mutation (N21K) in combination with R18A designed to replace the R18 ring with a new positively charged ring in the central pore (K21 ring). In contrast to all single R18 mutants tested previously^14^, R18A/N21K acquired second-site compensatory mutations in CA that restored its infectivity and replication upon propagation in MT4 cells (Figure 1). These mutations did not result in the introduction of any new charged rings in the central pore, indicating that the R18 ring can be functionally replaced by the K21 ring (Figure S2). Comparative structural and biochemical analysis of a rescued mutant, S16L/R18S/N21K (LSK), and a completely non-infectious mutant (S16N/R18S/N21K, NSK) revealed no differences in IP6 binding (Figure 2 A-C), in vitro CLP assembly (Figure 2D), or core formation in virions (Figure 3), suggesting that capsid assembly can occur at near-WT efficiency even in absence of the R18 ring and the molecule of IP6 with which interacts. In contrast, we observed a severe defect in the ability of NSK to undergo reverse transcription in cells (Figure 4 A-C). This finding combined with NSK’s sensitivity to DNase in our endogenous reverse transcription assay (Figure 4 D-F) and reduction in particles upon viral membrane permeabilization in our TIRF assay (Figure 5 suggests that NSK capsids are inherently unstable and break open shortly after fusion. This interpretation is supported by our confocal microscopy analyses that demonstrate that NSK capsids are defective in nuclear envelope docking and nuclear entry. Overall, our findings establish a role for the central pore in promoting the sustained stability of HIV-1 capsids that appears to be independent of its ability to bind IP6.

Our finding that HIV-1 can replicate in the absence of the R18 ring is noteworthy in light of early structural studies that described the R18 ring as the only binding site for IP6^10,11^. While several subsequent studies have confirmed that the K25 ring can also coordinate an IP6 molecule^15,19,21,41,42^, the R18 ring likely binds IP6 with greater affinity owing to the tighter packing of its side chains and its closer proximity to the outer surface of the capsid. Our LSK and NSK crystal structures demonstrate that the R18:IP6 interaction is lost upon replacement of the R18 ring with the K21 ring, suggesting that the R18 ring and its interaction with IP6 are not absolutely required for HIV-1 replication. We previously came to a similar conclusion regarding the importance of the K25 ring based on the ability of second-site compensatory mutations to restore infectivity and replication to K25A. However, a key difference in the relative importance of these two positively charged rings is that while the R18 ring can facilitate replication as the sole electropositive ring within the central pore, the K25 ring requires the introduction of an additional electropositive ring in the absence of R18. We speculate that this is due to lower affinity between IP6 and the K25 ring relative to the R18 ring, leading to a requirement for additional clustering of positively charged side chains to efficiently coordinate IP6.

Our finding that NSK capsids are inherently unstable but not assembly-defective indicates that the central pore plays a role in maintaining the stability of the HIV-1 capsid post-entry. Most reported central pore mutations that reduce particle infectivity are unable to efficiently assemble capsids during particle maturation. Notable exceptions are R18G and R18L, which are capable of assembling into normal cones, but also display aberrant capsid morphologies^14,17,20^. Given the importance of the central pore in binding IP6 and the ability of IP6 to stabilize capsids, our finding that mutations within the central pore can destabilize capsids seems intuitive. However, we were unable to detect notable differences in the ability of IP6 to bind or stabilize LSK and NSK CA hexamers (Figure 2 A and B), suggesting that the differences in the inherent stability of LSK and NSK capsids is unrelated to their ability to bind IP6. One caveat regarding this conclusion is that our crystallography and thermal stability measurements were performed with isolated, crosslinked CA hexamers. It is therefore possible that these experiments do not fully recapitulate the stabilizing effect of IP6 in the context of a complete capsid lattice composed of native hexamers and pentamers.

The LSK and NSK mutants differ at a single amino acid position (S16L vs. S16N). Why a leucine (LSK) at position 16 restores infectivity ∼30-fold, whereas an asparagine (NSK) completely blocks infectivity in the context of the R18S/N21K mutations is unclear. This is especially surprising because the Ser-to-Asn substitution is a more conservative amino acid change than Ser-to-Leu. The lack of even subtle differences in the hexamers of LSK vs NSK suggests that any structural changes associated with the different mutations are either not present in the cross-linked proteins used for study or occur between capsomers within the lattice. There may also be functional differences, for instance in dNTP import. The central pore is the channel through which dNTPs are putatively imported into the capsid to facilitate viral DNA synthesis. Importantly, the R18 and K25 rings have each been proposed to play a role in this process^16,17,43,44^. Notably, it is the polar side-chains at residue 16 that negatively affect infectivity (S16 or N16) whereas the non-polar leucine (L16) increases it. It is therefore tempting to speculate that the hydrophobic substitution at position 16 is needed to promote the import of nucleotides into the capsid interior and disfavor dissociation back out of the pore. Whilst there is no direct evidence for the role of position 16 in nucleotide import, NSK is clearly impaired for reverse transcription compared to LSK.

However, we did not observe substantial differences in the ability of ATP to bind or stabilize LSK and NSK CA hexamers (Figure S7). Consistent with the key role that R18 plays in dNTP binding to the central pore, both LSK and NSK exhibited a ∼10-fold reduction in ATP binding affinity (Figure S7). This reduction in ATP binding is less severe than the ∼100-fold reduction exhibited by R18G CA hexamers^17^, suggesting that the N21K mutation partially restores dNTP binding in the absence of the R18 ring. Because LSK remains capable of reverse transcribing its genome with near-WT efficiency, we propose that this reduction in nucleotide affinity is not sufficient to alter the kinetics of viral DNA synthesis. Importantly, the lack of differences in nucleotide binding displayed by LSK and NSK support our conclusions that capsid stability, rather than dNTP import, is the primary mechanism driving the differences in LSK and NSK infectivity.

In summary, our findings highlight the flexibility of the placement of the positively charged rings within the central pore of HIV-1 capsomers while simultaneously demonstrating the sensitivity of HIV-1 to changes within the central pore. Furthermore, our findings demonstrate the critical roles that the central pore plays in both the assembly of the HIV-1 capsid during particle maturation and sustained stability of the capsid post-entry.

## Materials and Methods

### Cells

MT4 and SupT1 T-cell lines were acquired from American Type Culture Collection (ATTC) and cultured in Roswell Park Memorial Institute (RPMI) 1640 medium supplemented with 10% fetal bovine serum, 2mM L-glutamine, penicillin (100U/mL), and streptomycin (100 μg/mL). HEK 293T, HeLa, and TZM-bl cells were acquired from ATCC or the NIH AIDS Reagent Program (TZM-bl) and cultured in Dulbecco’s modified Eagle’s medium (DMEM) supplemented as above. The previously described 293T Parental and IPPK KO cells^45^ were cultured in DMEM supplemented as above. Whole blood was acquired from healthy donors via the NCI-Frederick Research Donor Program, and PBMCs were isolated using a Ficoll gradient (Histopaque-1077; Sigma no. 10771). PBMCs were cultured in RPMI 1640 supplemented as above and stimulated with 2 μg/mL phytohemagglutinin P (PHA-P) for 5 days prior to infection. Culture medium was supplemented with 50 U/mL IL-2 during replication.

### Plasmids

pNL4-3, a lab-adapted, subtype B infectious molecular clone^46^, was used to transfect MT4 cells for replication and forced evolution experiments and used to transfect 293T cells to produce virus for infectivity assays. pNL4-3/KFS (env-)^47^ was used as a non-replicating control for replication and forced evolution experiments. pNL4-3/KFS was also used to transfect 293T along with a plasmid expressing VSV-G envelope to produce virus used to infect SupT1 cells for quantification of viral DNA synthesis. For confocal microscopy experiments, virions labeled with YFP-tagged HIV-1 Vif-resistant APOBEC3F mutant^48^ were prepared by co-transfection of 293T cells with pNL4-3/KFS containing WT or mutant CA, pA3F-E289K-YFP, and a VSV-G expression plasmid^49^. The A3F-E289K-YFP construct was generated by site-directed mutagenesis of pA3F-YFP^29^. An env-/vpr-, luciferase encoding pNL4-3-based plasmid (pNL4-3.Luc.R-.E-)^50^ was used to co-transfect 293T cells along with VSV-G to generate virus used for infectivity experiments in 293T Parental and IPPK KO target cells. HIV-1 CA mutant pNL4-3 plasmids were generated by subcloning DNA fragments purchased from Twist Bioscience. For cryo-ET and endogenous reverse transcription experiments, virions were produced in 293T cells via co-transfection of pMDG2, a packaging plasmid encoding VSV-G (Addgene #12259), pCRV GagPol^51^, and pCSGW^52^ as described previously. For TIRF experiments, virions were produced as above with a pCRV GagPol iEGFP plasmid^53^.

### Replication Kinetics

3 x 10^6^ MT4 cells were transfected with 3 μg pNL4-3 DNA in 0.7 mg/mL DEAE dextran dissolved in Dulbecco’s Phosphate Buffered Saline (DPBS) for 15 minutes at 37°C. The transfection mix was removed by centrifugation and the transfected cells were resuspended in complete RPMI medium. MT4 cultures were split at a 1:1 ratio and supernatant samples were collected every 2-4 days for up to three months. To determine the kinetics of viral replication, HIV-1 reverse transcriptase (RT) activity in the supernatant was quantified using a previously described ^32^P-based assay^54^. PBMCs were activated with PHA-P for 5 days and subsequently infected with 3 RT counts per minute (cpm) per cell for 2 hours at 37°C. PBMCs were then centrifuged and resuspended in complete RPMI medium supplemented with 50 U/mL IL-2 for ∼2-3 weeks. Supernatant samples were collected every 2 days and media were replenished without splitting the cells. Viral replication kinetics were determined as above.

### Forced-evolution experiments

MT4 cells were transfected with pNL4-3 harboring mutations coding for amino acid substitutions in the central pore of CA, and supernatants were collected every 2-4 days for up to three months. Cells and supernatant were collected at the peak of replication and genomic DNA was isolated using the QIAmp DNA Blood Mini kit (Qiagen – Catalog # 51106). Genomic DNA was subjected to polymerase chain reaction (PCR) using primers specific for the HIV-1 GagPol coding region. Amplified GagPol fragments were treated with ExoSAP-IT PCR Cleanup Reagent (Thermo Fisher Scientific Catalog # 78201.1.ML) and submitted for Sanger sequencing. RT activity in viral supernatants collected at the peak of replication was quantified, and 3 x 10^5^ RT counts per minute (cpm) of virus was used to inoculate 3 x 10^6^ fresh MT4 cells. Replication kinetics were measured to confirm reversion to wild-type replication, and genomic DNA isolation, PCR, and sequencing was repeated as above.

### Single-cycle infectivity

293T cells were transfected with pNL4-3 plasmids overnight using linear polyethylenimine (PEI) and media were changed the following day. Virus in the supernatant was collected at 24 hours post media change, filtered through a 0.45 μm syringe filter, and stored at −80°C. Virus was quantified by ^32^P RT assay and equal amounts of virus were used to infect TZM-bl cells. At 48 hours post infection, TZM-bl cells were lysed with britelite plus reagent (Revvity #6066766) and luminescence was quantified to determine infectivity. All measurements were normalized to signal acquired from WT-infected samples. For single-cycle infectivity measurements in IP6-depleted target cells, equal numbers of 293T Parental and IPPK KO cells were seeded in matched plates and infected with VSV-G-pseudotyped, luciferase-encoding, Δ1vprΔ1env (pNL4-3.Lu.R-.E-) virions. Luciferase was quantified as above to determine the infectivity of each virus in each cell line.

### Protein production and purification

The capsid (CA) proteins were expressed in *E.coli* C41 cells overnight at 25°C, lysed in lysis buffer (50 mM Tris-HCl (pH 8.0), 200 mM NaCl, 20% BugBuster, Protease inhibitor tablets, 1 mM DTT) and centrifuged (24,000 rpm, 1 hour). The supernatant was precipitated with 25% ammonium-sulphate (wt/vol) followed by centrifugation (13,000 rpm, 20 min). The precipitated CA was resuspended and dialysed against 50 mM MES (pH 6.0), 20 mM NaCl, 1mM DTT. The CA protein was further purified via a cation-exchange column with a gradient from 20mM-1M NaCl followed by size exclusion chromatography with Tris pH 8.0, 20 mM NaCl, 1mM DTT, concentrated and snap frozen.

The disulfide-stabilised CA hexamer was expressed in *E.coli* C41 cells, lysed with sonication in 50 mM Tris-HCl (pH 8.0), 200 mM NaCl, 20% BugBuster 20 mM 2-mercaptoethanol, Protease inhibitor tablets. Lysate was then cleared by centrifugation (24,000 rpm, 1 hour). The supernatant was precipitated with 25% ammonium-sulphate (wt/vol) followed by centrifugation (13,000 rpm, 20 min). The pelleted material was resuspended in 50 mM citric acid (pH 4.5) and dialysed against 50 mM Citric acid (pH 4.5), 20 mM 2-mercaptoethanol. The protein was further dialysed with multiple changes into 50 mM Tris-HCl (pH 8.0), 1 M NaCl, 20 mM 2-mercaptoethanol. Reducing agent was removed by dialysis against 50 mM Tris (pH 8.0), 1 M NaCl, and then finally into 20 mM Tris (pH 8.0), 40 mM NaCl. Assembled hexamers were identified by non-reducing SDS–PAGE and were further purified via anion-exchange (2X 5ml HiTrap® Q Fast Flow, GE Healthcare) in 20 mM Tris-HCl (pH8.0) 40 mM-1M NaCl gradient followed by size exclusion chromatography (Superdex 200) in 20 mM Tris (pH 8.0), 40 mM NaCl. Peak fractions were pooled and concentrated before snap freezing in 100µl aliquots and storage at −80°C.

### X-ray crystallography

CA hexamer protein was prepared exactly as described previously^11^.12 mg/ml hexamer was mixed with 1-5mM of IP6. Crystals were grown at 17 °C by sitting-drop vapour diffusion in which 100 nl protein was mixed with 100 nl precipitant and suspended above 80 µl precipitant in the MORPHEUS I screen^55^. Crystals were flash-cooled in liquid nitrogen and data was collected at beamline I24 at Diamond Light Source. The data sets were processed using the CCP4 Program suite^56^. Data were indexed and integrated with iMOSFLM and scaled and merged with AIMLESS^57^. Structures were solved by molecular replacement using the model 6ES8 in PHASER^58^ and refined using REFMAC5 ^59^. Between rounds of refinement, the model was manually checked and corrected against the corresponding electron-density maps in COOT^60^. Final figures were rendered in The PyMOL Molecular Graphics System, Version 1.5.0.4 Schrödinger, LLC.

### Nano-differential scanning fluorimetry

DSF measurements were performed using a Prometheus NT.48 (NanoTemper Technologies) over a temperature range of 20–95°C using a ramp rate of 2.5°C / min. CA hexamer samples were prepared at a final concentration of 7.5 µM hexamer in Intracellular buffer (110 mM Potassium gluconate, 25 mM KCl, 5 mM NaCl, 2 mM MgCl2, 10 mM HEPES). Ligands were added at 15 µM. DSF measurements were performed in triplicate.

### Fluorescence polarization anisotropy

Fluorescence anisotropy measurements were performed at 22 °C on a Cary Eclipse Fluorescence Spectrophotometer (Agilent). Fluorescein-labelled ATP was obtained from Perkin Elmer and used for saturation binding experiments at a concentration of 2 nM. Proteins were diluted to give a range of two-fold dilutions with highest final hexamer concentrations of 4 µM for mutants with ATP and 400nM hexamer for all other samples. Proteins were mixed with either FITC-ATP or FITC-HCB to give final probe concentrations of 2 nM nand 1 nM respectively. Proteins and probes were mixed at the indicated concentrations and allowed to equilibriate at room temp. Fluorescence polarization was measured using Pherastar FSX (optic module FP 485 520 520). Samples were measured in triplicate with at least two independent runs. Binding curves were fit using GraphPad Prism (GraphPad Software, Inc.).

### In vitro assembly assays

CA proteins were dialysed against 50 mM MES (pH 6.0), 40 mM NaCl, 1 mM DTT. CA proteins at a final concentration of 100 µM were mixed with IP6 at 25°C (final concentration 1.25 - 5 mM). The apparent increase in absorbance reflecting increased light scattering (OD_350_) was measured using a PHERAstar FSX Plate reader (BMG Labtech) in 384-well plate with shaking between each measurement at 25°C.

### Negative-stain electron microscopy

Samples were assembled at room temp with 200µM CA, 3mM IP6 for mutants and 400µM CA and 2.5mM IP6 for WT. 4 µl of sample from the assembly was put onto a glow discharged carbon coated grid (Cu, 400 mesh, Electron Microscopy Services), washed and stained with 2% Uranyl-acetate. Micrographs were taken at room temperature on a Tencai Spirit (FEI) operated at an accelerated voltage of 120 keV and Gatan 2k × 2 k CCD camera. Images were collected with a total dose of ∼20 e^-^/Å^2^ and a defocus of 1–3 µm. Cryo-electron Tomography

Virus-like particles were produced in HEK293T as previously described^14^. Supernatants were filtered through a 0.45µm and 0.22μm filter. Particles were concentrated by ultracentrifugation over a 20 % (wt/vol) sucrose cushion (2 hours at 28,000 rpm in a Beckman SW32 rotor; Beckman Coulter Life Sciences). The pellet was resuspended in PBS. Grids were prepared by the plunge-freeze method in liquid ethane using a FEI Vitrobot Mark II. Data collection and analysis was performed as previously described^14^ but briefly: Tomographic tilt series of LSK and NSK were acquired between −60° and +60° with increments of 3° using a dose symmetric scheme using Serial-EM^61^. Tomograms were reconstructed using IMOD 4.9^62^. The alignment of 2D projection images of the tilt series was performed using gold beads as fiducial markers, tomograms were reconstructed by back projection.

### Reverse transcription in cells

2.5 x 10^6^ 293T cells were seeded into 10 cm dishes, cultured overnight, and co-transfected with pNL4-3 and a VSV-G expression vector. Virus-containing supernatant was collected and filtered as above and treated with Benzonase nuclease (Millipore Sigma #E1014) to remove carryover plasmid DNA. Virus was then pelleted through an 8% OptiPrep cushion and resuspended in complete DMEM containing amplification grade DNase I and 10 mg/mL MgCl_2_. Virus was aliquoted, quantified by ^32^P RT assay, and stored at −80°C until use. 5 x 10^5^ SupT1 cells per timepoint were infected with 10 RT cpm per cell of each virus for 2 hours at 4°C. Cells were harvested at 1, 2, 3, 6, 9, and 12 hours post-infection, washed once with PBS, lysed, and frozen for subsequent processing. DNA was isolated using the QiaAMP DNA Blood Mini kit according to the manufacturer’s instructions. Viral transcripts were quantified by qPCR of 5 μL of extracted DNA with 10 μL Taqman Fast Advanced Master Mix (Applied Biosystems #4444557) and 5 μL primer:probe mix on a QuantStudio 3 Real Time PCR system (Applied Biosystems). Previously described primers pairs and probes were used to detect early (minus-strand strong stop; MSSS) and late (second-strand transfer; SST) RT products and 2-LTR circles^63^. CCR5 DNA was also quantified to control for cell number. Probes were labeled at the 5’ end with 6-carboxyfluorescein (FAM).

### Endogenous reverse transcription

For ERT reactions HEK293T cells were seeded on 10cm dishes and transfected as above but without VSVG-expressing plasmid. We used four dishes per virus. After 6h cells were washed with PBS to remove excess DNA and then incubated for 3 days in DMEM with 2% FBS. Supernatants were filtered through 0.45μm filters and incubated with 1/1000 benzonase (70746, Millipore) at room temperature for 1h. Supernatants were then spun through 20% sucrose cushion for 2h at 28k in SW28 and resuspended on ice with 500ul 100mM Tris pH 8, 150mM NaCl. Aliquots were frozen for further use. ERT reactions were set up in 96-well plates in 20μl volumes. 10μl of freshly thawed VLPs were mixed with buffer containing 100mM Tris pH8, 150mM NaCl, 2mM MgCl_2_, 0.5mM TCEP and 1mg/ml of BSA (B8667, Sigma). Reactions were supplemented with desired concentrations on IP6 (Sigma, 593648), dNTPs (28-4065-51, GE Healthcare), DNase (Sigma, DN25) at final concentration 0.1mg/ml and 50nM.

Streptolysin O (SLO, Sigma, S5265). Plate was then covered with adhesive film, spun down and incubated at 37°C for 5h. To purify DNA we used Blood and Tissue Qiagen kit as follows: add 20μl reaction volume to 180μl PBS and then mix with 200μl lysis buffer, continue as per manufacturer instructions and elute DNA in 40μl of water. DNA was quantified using Power SYBRGreen (Applied Biosystems) ABI StepOnePlus PCR System using standard SYBRGreen protocol. For standards we used CSGW vector that was used to generate VLPs. Primers were for early products (MSSS: minus strand strong stop), intermediate products (FST: forward strand transfer) and late products (SST: second strand transfer). Reactions were run as duplicates.

### Streptolysin O (SLO) production

The 54kDa (residues 101-571) version of the SLO was tagged with 6xHis by cloning into pOPTH-Tev vector for production in *E.coli* C41 strain. Cells were lysed with 20% BugBuster in 25mM Tris pH8, 150mm NaCl, 1mM DTT, benzonase and lysozyme. Lysate was loaded onto a Ni column in 25mM Tris pH8, 150mM NaCl, 1mM DTT, 10mM Imidazole, washed with buffer with 10mM Imidazole, followed by 40mM Imidazole and then eluded with 200mM Imidazole. 1ml fractions were then checked on a polyacrylamide gel, peak fractions combined and dialysed to 25mM Tris ph8, 150mM NaCl and 1mM DTT. Protein was then aliquoted and stored at −70°C.

### TIRF microscopy

TIRF microscopy was carried out as previously described^14^. pCRV GagPol iEGFP construct was first generated by inserting EGFP between MA and CA of Gag with EGFP sequence flanked by protease cleavage sites for EGFP release from Gag upon maturation^53^. To generate virions, constructs with corresponding capsid mutations were transfected at a ratio of 1 to 4 of pCRV GagPol iEGFP to pCRV GagPol. 6 hours after transfection, cells were washed and then incubated for 3 days with 2% FBS containing media. Virus containing supernatants were filtered as above and quantified by RT activity. Supernatants were diluted in PBS to equalize to RT units across viruses then placed on 8-well glass Ibidi dishes precoated with poly-L-lysine (Sigma, P4707) for binding. After two PBS washes, viruses were incubated for 30 min with 20nM SLO in the presence or absence of 50μM IP6 in in reaction buffer (SRB, 100mM Tris-HCl pH 8, 150mM NaCl). Untreated wells were used as control for normalisation. After incubation wells were washed 2 times with PBS then fixed with 4% formaldehyde for 20 min. Images were acquired with a Nikon TIRF inverted microscope with a 100x/1.49NA oil-immersion objective, a 1.5x intermediate magnification and Prime95B sCMOS camera from Photometrics resulting in a 74nm pixel size. Particle numbers were analysed in Fiji, where images were median filtered, and background subtracted. An intensity threshold was used to create a mask and a watershed step allowed separation of touching particles. ROI were filtered by area within 1 to 200 pixels.

### Quantifying nuclear envelope docking and nuclear entry

Virions labeled with A3F-E289K-YFP were prepared by co-transfection of 293T cells with pNL4-3/KFS encoding WT or mutant CA, pA3F-E289K-YFP, and VSV-G expression vector^28^. Supernatants from the transfected cells were collected 24 hours post-transfection, clarified using a 0.45 µm membrane filter, and concentrated by ultracentrifugation (100,000 × g) for 1.5 hours at 4°C through a 20% sucrose cushion (wt/vol) in 1X Dulbecco’s Phosphate-Buffered Saline with calcium and magnesium (PBS). The concentrated virus was resuspended in 500 µl DMEM and the number of YFP-labeled virions was quantified by single virion analysis^64^. To quantify nuclear envelope association and nuclear import of HIV-1, HeLa-based cell lines were seeded in ibiTreated μ-slides (3 × 10⁴ cells/well; Ibidi) one day prior to infection. Cells were challenged with an equal number of A3F-E289K-YFP-labeled virus via spinoculation (1,200 × *g* for 1 hour at 15°C) in the presence of polybrene (10 µg/ml). Following spinoculation, the medium was replaced with prewarmed medium containing BioTracker 650 Red Nuclear Dye (0.5 µl/ml; Millipore). At 6 hours post-infection, the cells were fixed with 4% paraformaldehyde in 1X PBS for 10 minutes. Confocal z-slices at the equatorial plane of the cells were acquired using a Nikon Eclipse Ti-E microscope equipped with a Yokogawa CSU-X1 spinning disk unit and a Plan-Apochromat 100× N.A. 1.49 oil objective, using 514-nm (YFP) and 647-nm (BioTracker650 dye) lasers for illumination. Images were captured using a TwinCam system (Cairn) equipped with a 565-nm splitter and two iXon Ultra (Andor) cameras and analyzed with Nikon Elements or ImageJ^65^. Each field of view (88 µm²) contained an average of 8 cells, and 7 fields were acquired per sample. A custom MATLAB program^1^ was used to quantify the efficiency of nuclear envelope association and nuclear import of YFP signals. First, YFP signals were detected using Localize^66^. Second, nuclear envelope and nuclear masks were generated from the BioTracker 650 signal. Finally, colocalization of each YFP spot with the nuclear envelope and nuclear masks was assessed. The percentage of cytoplasmic YFP spots colocalized with nuclear envelope masks and the percentage of total YFP spots colocalized with the nuclear masks was calculated. For display purposes, a pixel-averaging filter was applied to the images, and contrast was adjusted.

### Statistical Analysis

All experiments were repeated at least three times unless otherwise stated. The number of repeats (N), determination of error, and statistical analysis is listed for each specific experiment in the figure legends.

**Figure S1. K25A acquires N21K upon propagation in MT4 cells.** (A) MT4 cells were transfected with WT or CA-K25A infectious molecular clones (NL4-3) and viral replication was monitored by RT assay. (B) WT or K25A virus collected from the peak of replication in (A) were used to infect MT4 cells and viral replication was monitored by RT assay.

**Table S1. CA mutations acquired by central pore mutants subjected to forced evolution.** MT4 cells were transfected with infectious molecular clones harboring the indicated mutations. Virus collected from the peak of replication was used to infect fresh MT4 cells, and the process was repeated until replication kinetics resembled WT. Compensatory mutations acquired upon propagation were identified by Sanger sequencing of GagPol DNA fragments amplified by PCR from genomic DNA isolated from cells at the peak of viral replication. HIV-1 mutant clones used to initiate each forced evolution experiment in this study are listed along with the number of passages and compensatory mutations identified in CA.

**Figure S2. Locations of compensatory mutations acquired upon propagation of R18A/N21K and R18S/N21K.** (A) Cross section of the central pore of the CA hexamer (based on PDB 6R6Q) showing the locations of H12 (orange), S16 (pink), and N21 (magenta) relative to the R18 and K25 IP6-binding rings. (B) 90° rotation of (A) to depict the location of A31T (red) on the luminal surface of the CA hexamer.

**Figure S3. A serine at position 18 facilitates enhanced rescue by compensatory mutations.** Single-cycle infectivity of HIV-1 CA mutants harboring the R18A/N21K changes (A) or the R18S/N21K (B) changes with compensatory mutations identified in this study quantified by luminescence in infected TZM-bl cells. Virus stocks were normalized by RT assay prior to infection of TZM-bl cells. The R18A/N21K data presented for comparison in (A) are the same that is presented in Figure 1B. The SK, LSK, and SKT data presented for comparison in (B) are the same that is presented in Figure 1H. Error bars depict the mean ± s.e.m. from at least 3 independent measurements. Abbreviations: R18S/N21K (SK); S16L/R18S/N21K (LSK); R18S/N21K/A31T (SKT).

**Figure S4. Compensatory mutations restore replication in MT4 cells.** Representative spreading infection experiment depicting the replication kinetics of WT HIV-1 or defective (R18A, R18S, R18A/N21K, SK, NSK, LSR) and rescued (LSK, SKT, SKI) central pore mutants in MT4 cells. Spreading infection experiments were initiated by transfection of MT4 cells with infectious molecular clones and replication kinetics were monitored by RT assay. The Env(-) clone pNL4-3/KFS is included as a negative control. Abbreviations: R18S/N21K (SK); S16L/R18S/N21K (LSK); R18S/N21K/A31T (SKT); R18S/N21K/T216I (SKI); S16N/R18S/N21K (NSK); S16L/R18S/N21R (LSR).

**Figure S5. Compensatory mutations partially restore replication to R18S/N21K (SK) in primary peripheral blood mononuclear cells (PBMCs)**. Cells isolated from whole blood collected from 4 individual donors were activated for 5 days with PHA-P and infected with RT-normalized WT, SK, LSK, SKT, or SKI. Replication kinetics were monitored by RT assay. Abbreviations: R18S/N21K (SK); S16L/R18S/N21K (LSK); R18S/N21K/A31T (SKT); R18S/N21K/T216I (SKI).

**Figure S6. LSK and NSK CA assembles less efficiently than WT CA at low IP6 concentrations.** In vitro assembly kinetics of 100 μM of the indicated recombinant mature CA protein in the presence of 1.5 mM (A) and 3 mM (B) IP6, as determined by measuring absorbance at 350 nm over time. Abbreviations: S16L/R18S/N21K (LSK); S16N/R18S/N21K (NSK).

**Figure S7. LSK and NSK crosslinked CA hexamers bind weakly to ATP and dATP.** (A) Thermal stability measurements for the indicated crosslinked CA hexamers alone or in the presence of ATP or dATP, as measured by differential scanning fluorimetry. Error bars depict the mean ± s.e.m. from at least 3 independent experiments. (B) ATP binding kinetics for the indicated crosslinked CA hexamers, as measured by fluorescence polarization. Error bars depict the mean ± s.e.m. from at least 3 independent experiments. Abbreviations: S16L/R18S/N21K (LSK); S16N/R18S/N21K (NSK).

**Figure S8. Rescued central pore mutants display mild IP6 dependency in target cells.** VSV-G pseudotyped, luciferase-encoding virions were used to infect 293T parental and IPPK KO target cells. Specific infectivity was measured at 48 hours post-infection by quantifying luminescence after lysis of infected cells. Infectivity for each virus in IPPK KO target cells is expressed as a percentage of its infectivity in parental cells. Error bars depict the mean ± s.e.m. from 8 independent experiments. Statistical significance was determined by unpaired Student’s t-test (p-value summary: >0.05 = not significant; <0.05 = *; <0.01 = **; <0.001 = ***; <0.0001 = ****. Abbreviations: R18S/N21K (SK); S16L/R18S/N21K (LSK); R18S/N21K/A31T (SKT); R18S/N21K/T216I (SKI).

## Acknowledgements

Research in the Freed and Pathak laboratories is supported by the Intramural Research Program of the Center for Cancer Research, National Cancer Institute, National Institutes of Health. ABK was supported in part by an Intramural AIDS Research Fellowship and the National Institute of Allergy and Infectious Diseases (K99AI174891-01). EOF would like to acknowledge our collaborative interactions with the Pittsburgh Center for HIV Protein Interactions (U54AI170791); EOF and VKP would like to acknowledge our collaborative interactions with the Behavior of HIV in Viral Environments Center (U54AI170855). Research in the James lab is supported by the MRC (UK; U105181010), a Wellcome Trust Investigator Award (200594/Z/16/Z), and a Wellcome Trust Collaborator Award (214344/A/18/Z). The content of this publication does not necessarily reflect the views or policies of the Department of Health and Human Services, nor does mention of trade names, commercial products, or organizations imply endorsement by the U.S. Government.

## Author Contributions

**Alex B. Kleinpeter**: Conceptualization, Investigation, Formal Analysis, Visualization, Writing-Original draft preparation, Writing-Reviewing and Editing. **Donna L. Mallery**: Investigation, Formal Analysis, Visualization. **Anna Albecka**: Investigation, Formal Analysis, Visualization. **Ryan C. Burdick**: Investigation, Formal Analysis, Visualization, Writing-Reviewing and Editing. **Nadine Renner**: Investigation, Formal Analysis, Visualization. **J. Ole Klarhof**: Investigation, Formal Analysis, Visualization. **Boglarka Vamos**: Investigation, Formal Analysis, Visualization. **Vinay K. Pathak**: Supervision, Writing-Reviewing and Editing. **Leo C. James**: Conceptualization, Supervision, Writing-Reviewing and Editing. **Eric O. Freed**: Conceptualization, Supervision, Writing-Reviewing and Editing.

