## Supplementary figures and images for "The central pore of HIV-1 capsomers promotes sustained stability of the viral capsid"

### Supplementary Figure 1

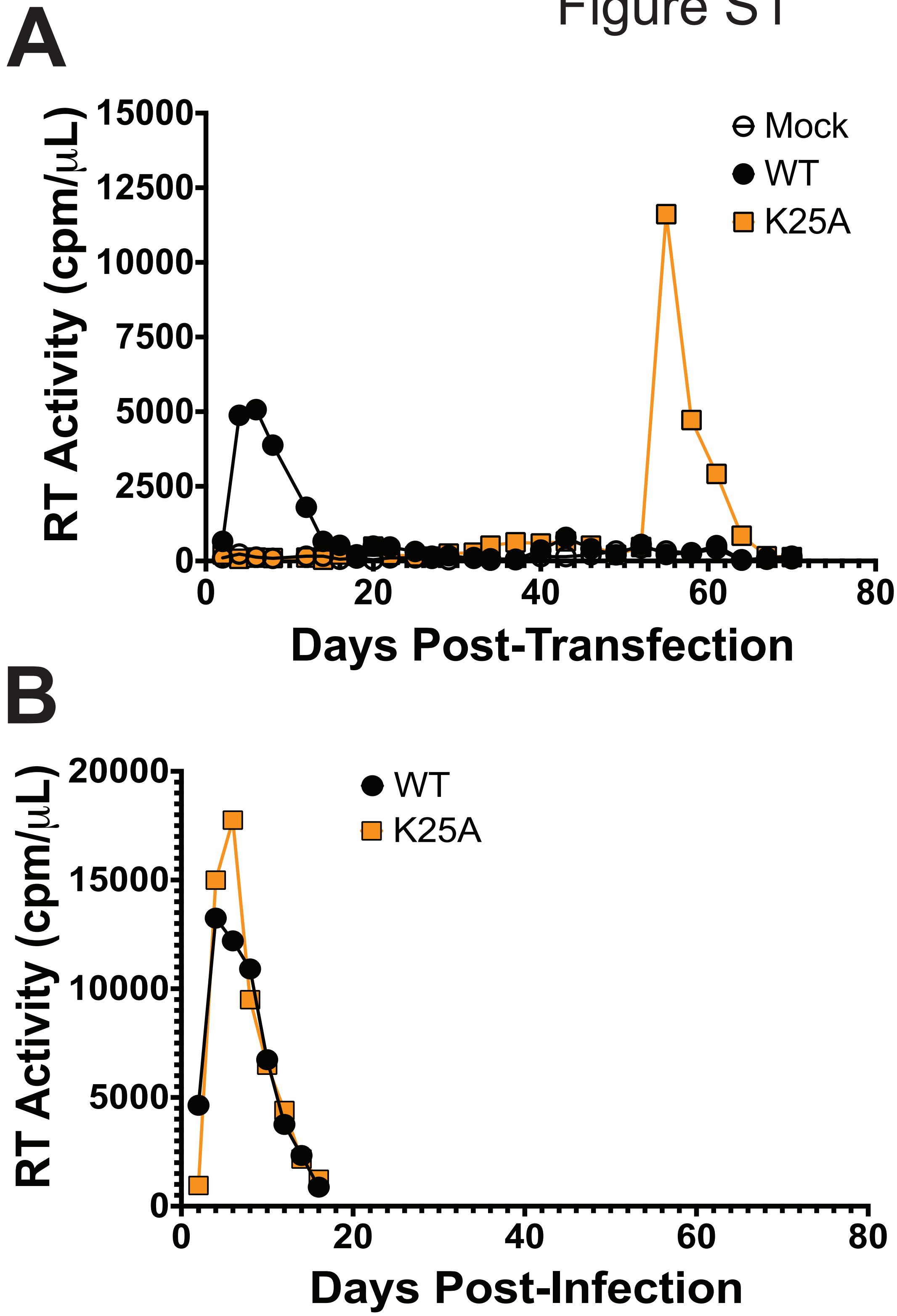

### Supplementary Figure 2

Figure S2

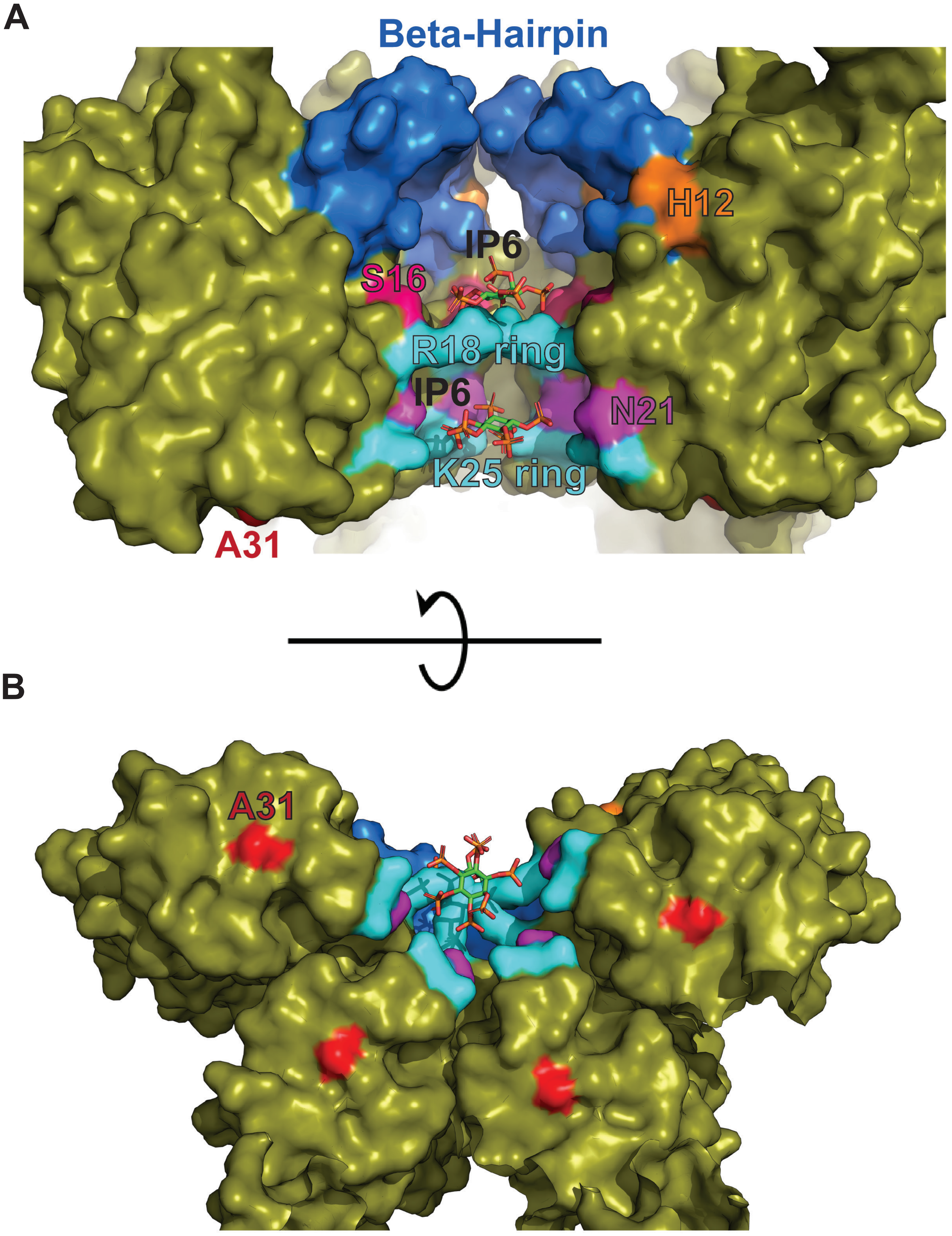

### Supplementary Figure 3

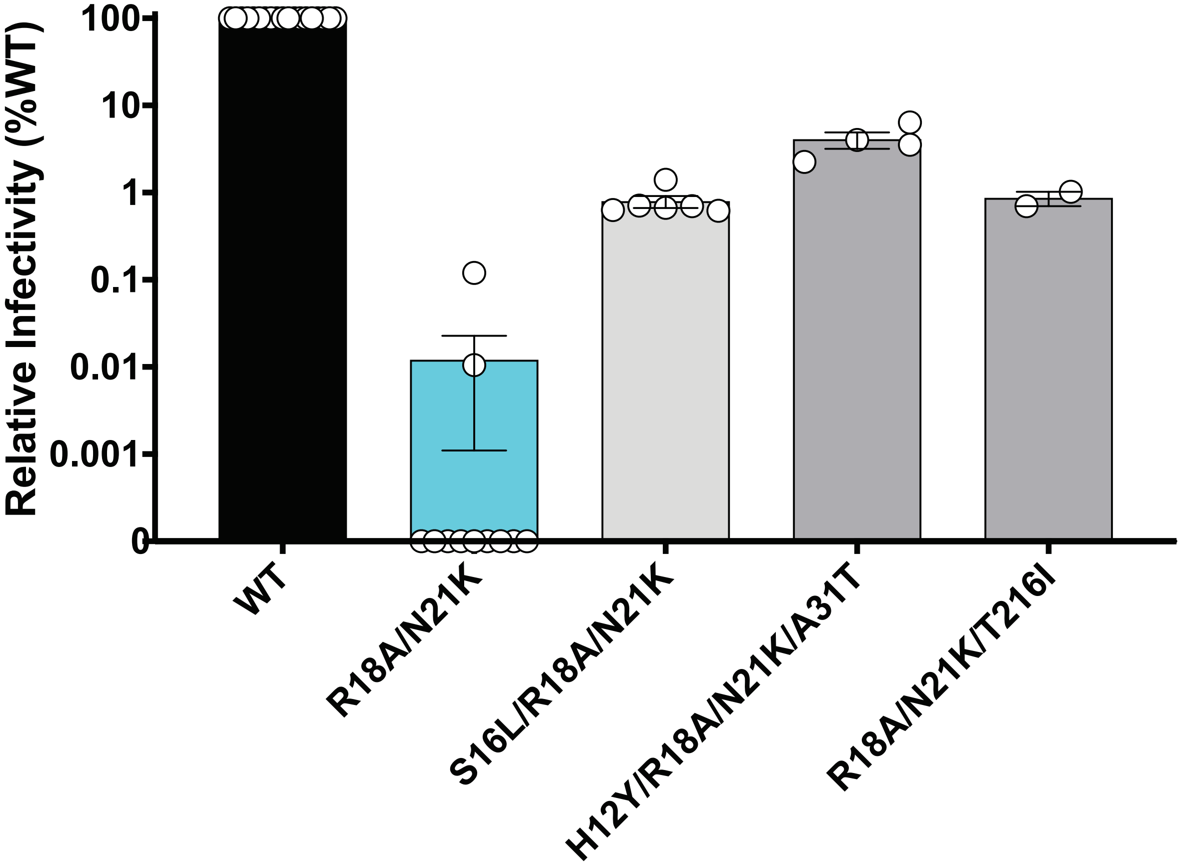

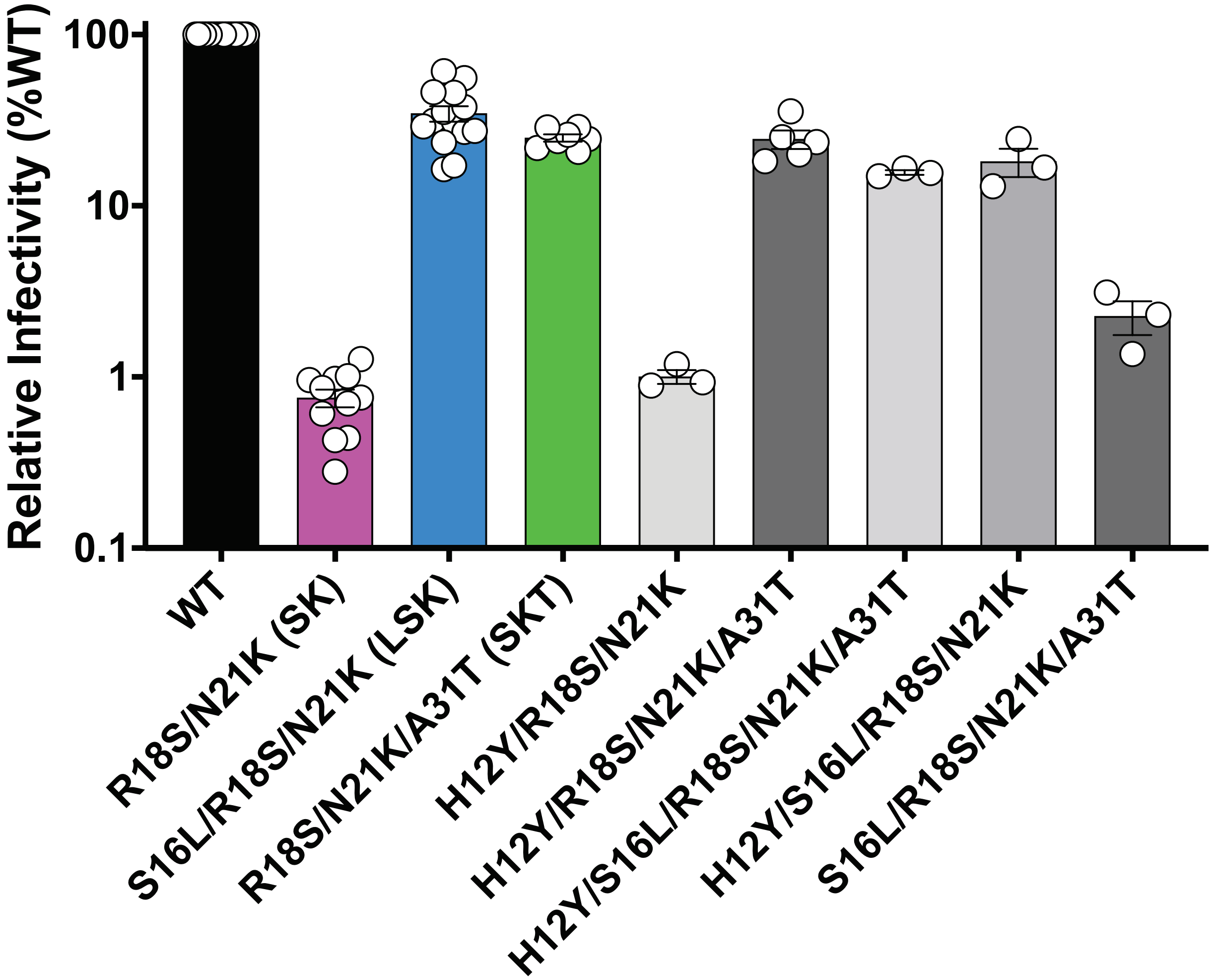

### Supplementary Figure 4

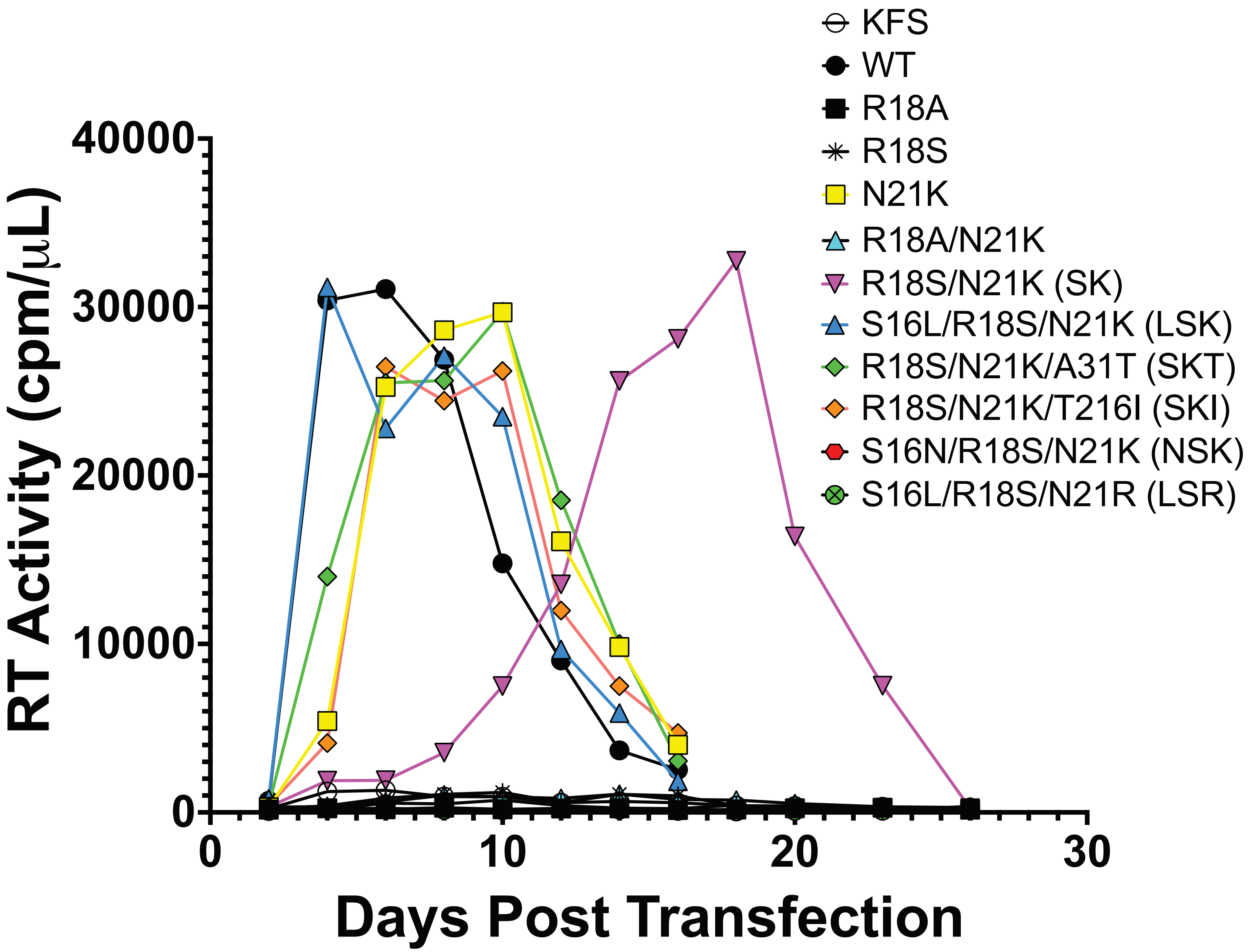

### Supplementary Figure 5

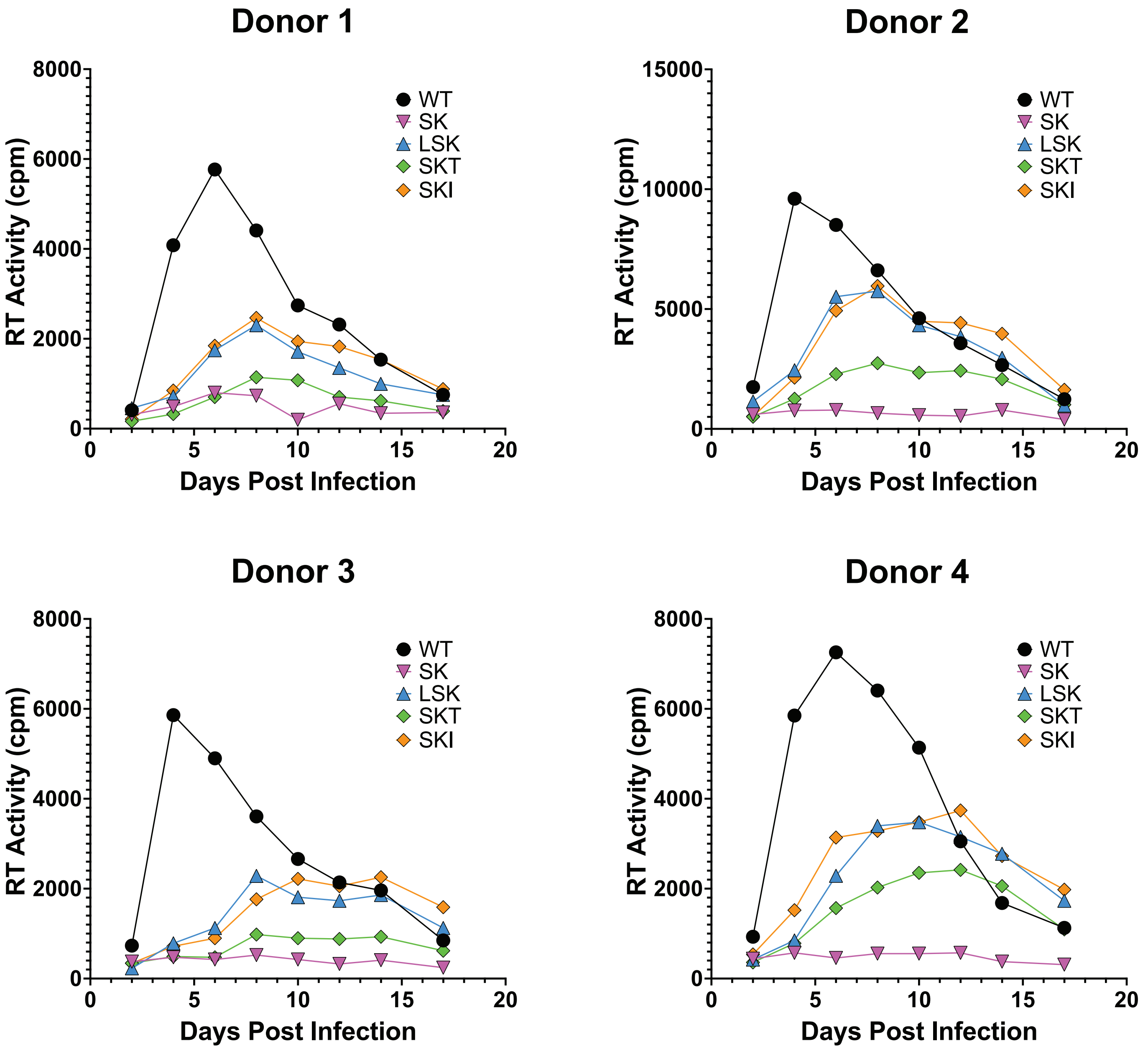

### Supplementary Figure 6

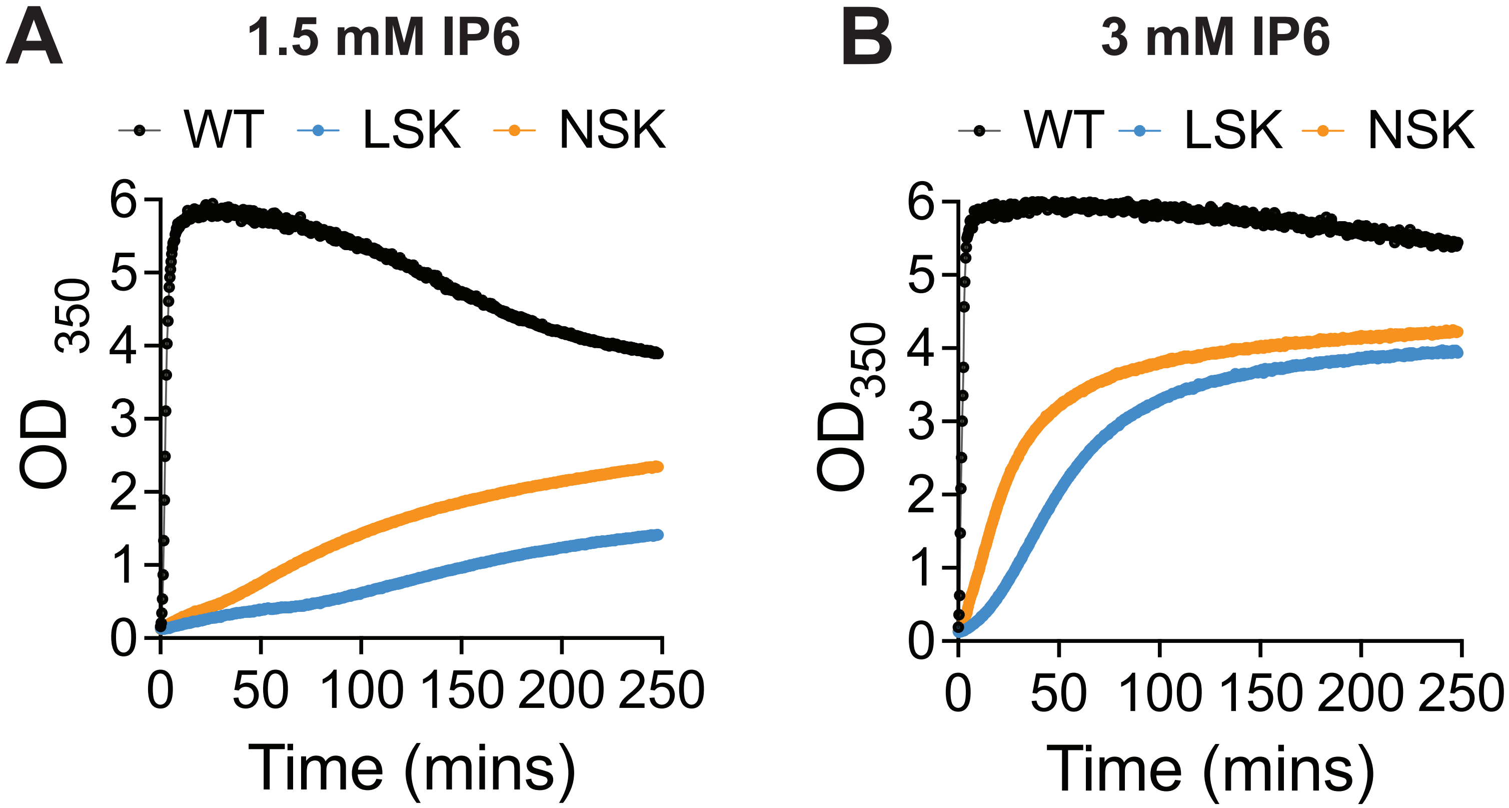

### Supplementary Figure 7

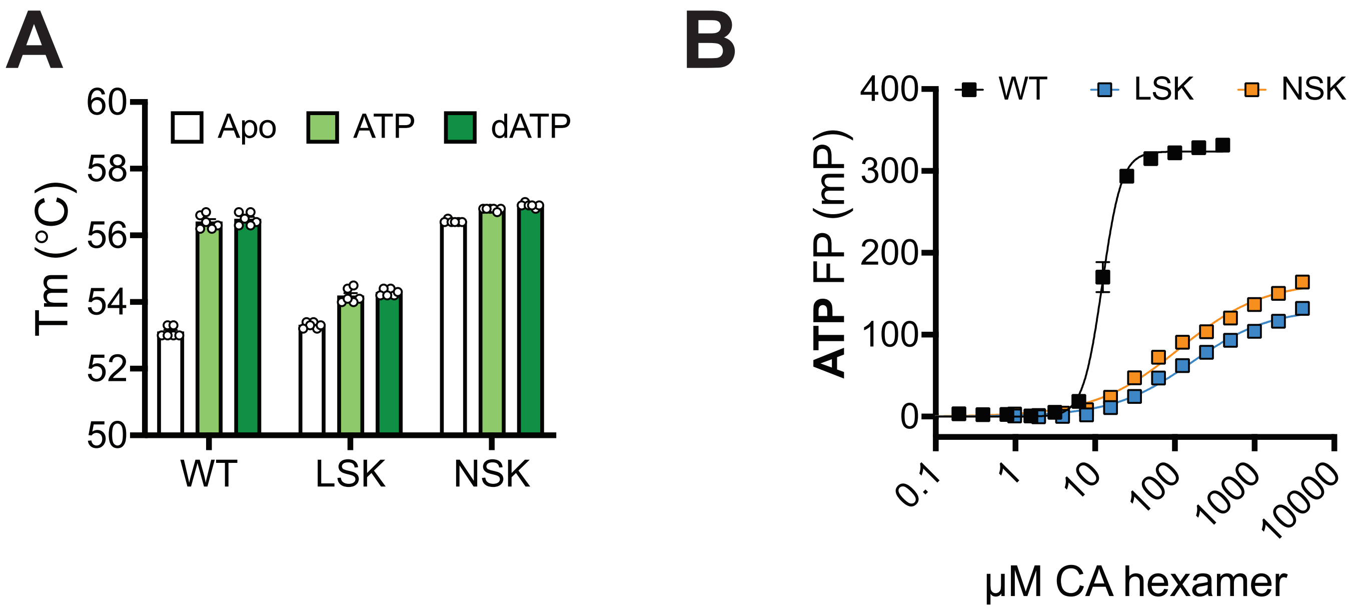

### Supplementary Figure 8

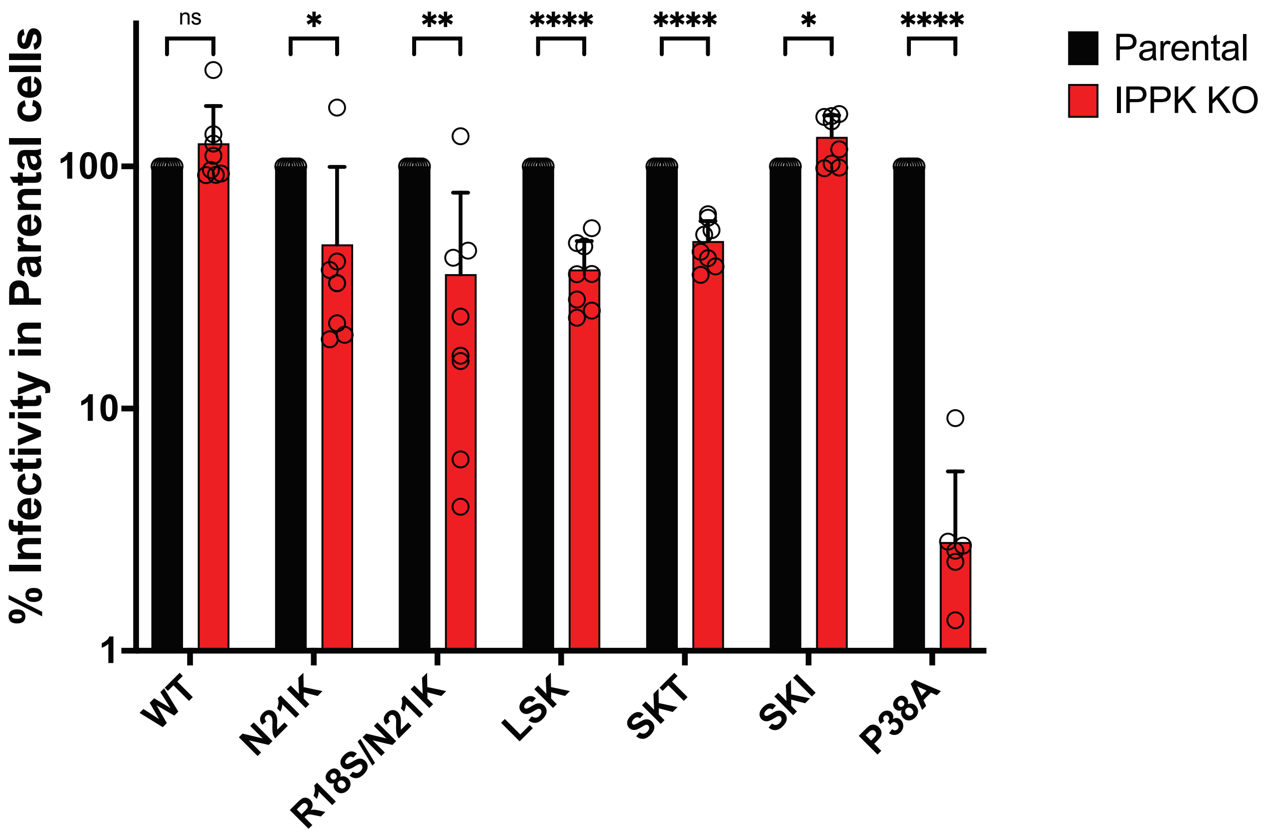
