## Supplementary Table 1 for "The central pore of HIV-1 capsomers promotes sustained stability of the viral capsid"

### Table S1

| Transfected Clone | Passages | CA Mutations Acquired |
|---|---|---|
| K25A | 2 | N21K |
| R18A/N21K | 3 | S16L, A18S |
| R18A/N21K | 3 | H12Y, A31T |
| R18A/N21K | 3 | L56V, E187K |
| R18S/N21K | 2 | T216I |
| R18S/N21K | 2 | T216I |
| R18S/N21K | 2 | A31T |
| R18S/N21K | 2 | I15M |
| S16N/R18S/N21K | 2 | N16L |
| S16L/R18S/N21R | 2 | R21K |
